# Orbitofrontal-hypothalamic projections are disrupted in hypermetabolic murine ALS model and human patients

**DOI:** 10.1101/2020.11.28.402065

**Authors:** David Bayer, Stefano Antonucci, Hans-Peter Müller, Luc Dupuis, Tobias Boeckers, Albert Ludolph, Jan Kassubek, Francesco Roselli

## Abstract

Increased catabolism is a new clinical manifestation of Amyotrophic Lateral Sclerosis. A dysfunction of lateral hypothalamus may drive hypermetabolism in ALS; however, Its causes and anatomical substrates are unknown. We hypothesize that disruption cortico-hypothalamic circuits may impair energy homeostasis in ALS. We used rAAV2 for large-scale projection mapping and image analysis pipeline based on Wholebrain and Ilastik to quantify projections from the forebrain to the latera hypothalamus of the SOD1(G93A) ALS mouse model as well as of the Fus^ΔNLS^ ALS mouse model. Expanded projections from agranular Insula, ventrolateral orbitofrontal and secondary motor cortex to lateral hypothalamus were found in two independent cohorts of the hypermetabolic SOD1(G93A) ALS model. The non-hypermetabolic Fus^ΔNLS^ ALS mouse model display a loss of projections from motor cortex but no change in projections from insula and orbitofronal cortex. 3T DTI-MRI data on 83 ALS patients and 65 controls confirmed the disruption of the orbitofrontal-hypothalamic tract in ALS patients. Converging murine and human data demonstrate the selective disruption of hypothalamic inputs in ALS as a factor contributing to the origin of hypermetabolism.

**Significance statement:** We provide a circuit perspective of the recently identified and medically relevant hyper-metabolic phenotype of Amyotrophic Lateral Sclerosis. We demonstrate the selective involvement of orbitofrontal, insular and motor cortex projections to hypothalamus in murine ALS models and in human patients. The enhanced pipeline for large-scale registration, segmentation projections mapping, the identification of new circuits target of neurodegeneration, and the relevance of these circuits in metabolic disturbances make this work relevant not only for the investigation of ALS but also for other neurodegenerative disease as well as for all conditions characterized by systemic energy imbalances.

## Introduction

Amyotrophic lateral sclerosis (ALS) is traditionally conceptualized as a neurodegenerative condition primarily affecting upper motoneurons, located in primary motor cortex, and lower motoneurons, located in spinal cord, whose dysfunction and loss determines a relentless, progressive and ultimately fatal motor impairment (Hardiman et al., 2017). More recently, hypermetabolism has been recognized as an additional, non-motor clinical feature of ALS (Dupuis et al., 2011). Epidemiological surveys have revealed that ALS patients display increased catabolism (Dupuis et al., 2011; Peter et al., 2017) and pointed out lower body-mass index constitutes a risk factor for ALS (Gallo et al., 2013). Furthermore, reduced levels of metabolic rate proxies such as plasma lipids and body fat content are predictors of survival of ALS patients (Dupuis et al., 2008; Dorst et al., 2011; Lindauer et al., 2013), i.e. weight loss is strongly correlated with shorter survival (Marin et al., 2011). Increased catabolism appears to be due to intrinsic hypermetabolism both in ALS patients (Steyn et al., 2018; Jesus et al., 2018; Ahmed et al., 2016; Desport et al., 2005; Bouteloup et al., 2009) as well as in mutant SOD1 ALS mice (Dupuis et al., 2004). Notably, metabolism and energy balance actually constitute promising targets for intervention, since increasing caloric intake has beneficial effects on survival, particularly on fast-progressing ALS patients (Ludolph et al., 2020).

Mechanistic insights into ALS hypermetabolism remain limited. Hypothalamic nuclei receive and integrate inputs coming from a large fraction of the brain (Gonzalez et al., 2016; Barbier et al., 2020a; Barbier et al., 2020b; Murata et al., 2019) in order to establish the proper balance of feeding, energy storage and energy expenditure (Berthoud, 2002). Thus, disruption of these inputs may be sufficient to drive metabolic imbalances. In fact, the overall architecture between large-scale networks appear to be disturbed in ALS (Commisso et al., 2018; Agosta et al., 2013; Schulthess et al., 2016; Heimrath et al., 2014; Dukic et al., 2019) and ALS-related pathobiochemistry affects a significant proportion of cortical and subcortical structures, in a pattern evolving with disease progression (Brettschneider et al., 2013; Braak et al., 2013). We have hypothesized that a similar degree of disruption and remodeling may take place in large-scale networks providing inputs to the feeding-regulating Lateral Hypothalamic Area (LHA) and that such a disruption may coincide with the appearance of the hypermetabolic phenotype. We have now identified the selective disruption of projections from insular and orbitofrontal cortex to LHA in ALS mouse model as well as in human ALS patients by a combination of retrograde rAAV-2 tracing and MRI-DTI tract tracing.

## Materials and Methods

### Animals

All experimental procedures involving animals were performed in agreement with the guidelines for the welfare of experimental animals issued by the Federal Government of Germany; all experiments were approved by the Regierungspräsidium Tübingen under the animal license- number 1390 and by the Ulm University Tierforschungszentrum committee. No effort was spared to implement “3R” guidelines for animal experimentation.

We obtained from Jackson Laboratories the following strains: B6SJL73 Tg(SOD1*G93A)1Gur/J (high-copy), (henceforth mSOD) and B6.Cg-Gt(ROSA)26Sor^tm6/(CAG74ZsGreen)Hze/J^ (henceforth ZsGreen). To generate the mSOD/ZsGreen double transgenic mice, hemizygous mSOD males were crossed with homozygous ZsGreen^+/+^ females as previously reported (Commisso et al., 2018); the progeny included mSOD^+^/ZsGreen^+/−^ mice at the expected mendelian rate. mSOD^+^/ZsGreen^+/−^ are henceforth going to be labelled mSOD1 and mSOD^−^/ZsGreen^+/−^will be labelled WT.

The heterozygous B6.Fus^ΔNLS/+^ (henceforth Fus) were provided by Luc Dupuis from the Faculté de Médecine, Strasbourg, France (Scekic-Zahirovic et al., 2016). For the generation of Fus/ZsGreen double transgenic mice, heterozygous Fus males were crossed with homozygous ZsGreen^+/+^ females. The progeny included Fus^−^/ZsGreen^+/−^ mice at the expected mendelian rate. Fus^−^/ZsGreen^+/−^ are henceforth going to be labelled kiFUS.

Mice were housed at 2 - 5 animals per cage, with unlimited access to food and water, a light/dark cycle of 12/12 hr and humidity between 40% and 60%. All mice expressing mSOD were routinely tested for motor impairment and euthanized in case of overt motor disability.

Since male and female mice have differ substantially in progression rates of clinical and biological manifestation of motoneuron disease (e.g., Ouali-Alami et al., 2018), the present study focused on male mice only.

### Viral vectors

Retrograde rAAV2 (Tervo *et al.*, 2016) vectors encoding for the pmSyn1-EBFP-Cre (addgene plasmid # 51507; kindly donated by Hongkui Zeng, Madisen *et al.*, 2015),were obtained from Addgene.

### Intracerebral injection

Intracerebral injections of rAAVs were performed as previously reported (Commisso *et al.*, 2018). Briefly, 1μl of viral suspension with a titer of 3×10^12^ vg/ml and 1 μl of 1% of Fast Green were freshly mixed and 300 nl of the mixture was loaded into a pulled-glass capillary. Pulling parameters were optimized to obtain a long and tapering capillary tip. Mice were pre-treated with buprenorphine (0.1 mg/kg) and meloxicam (1 mg/kg) 20 min before being anesthetized with 5% sevoflurane /95% O_2_; mice were then positioned into a stereotactic frame (David Kopf) and kept under continuous anesthesia with 3% sevoflurane/97% O_2_. Mice were positioned on a heated pad connected to a closed-loop system to maintain body temperature, monitored by a rectal probe, at 37°C. Upon incision of the scalp, a burr hole was prepared using a hand-held microdrive aiming the LHA at AP = −1.20, ML = −1.25, DV = −4.70 (according to Paxinos atlas, 2nd edition (Paxionos & Keith, 2001) refined for the mouse strain under consideration of the age). Under visual control, the pulled-glass capillary (tip closed) was moved into the drilled hole and gently pushed down to verify that meninges could be penetrated without deflection of the capillary. Thereafter, the pulled-glass capillary was withdrawn and opened at the very tip by gently touching with a microscissor. Before lowering the pulled-glass capillary to the final position inside the brain, a thin layer of DPBS^+Ca/+Mg^ was applied, generating a virus-free tip by capillary force. Since brain tissue might absorb some of the liquid out of the pulled-glass capillary by its own tissue/capillary force, during further movement, the pulled-glass capillary was lowered not slower than 2 mm/s. The viral suspension was injected using a Picosprizer microfluidic device; injection was performed during 5 min. The duration of one pulse was 10 ms; since the opening diameter of each capillary could be slightly different, the injection pressure was adapted (between 20 and 60 psi) to ensure a constant overall flow rate of 50-60 nl/min. After injection, the capillary was kept in place for 15 min and then removed with a continuous movement (1 mm/s). Remaining pressure in the capillary was released before withdrawing it to prevent any residual virus suspension to be discharged into the capillary thread.

### Immunohistochemistry

Mice (mSOD) were sacrificed at P (post-natal day) 40 (injection at P25) or at P110 (injection at P95); Fus mice were injected at P255 and sacrificed at P270.

Mice were trans-cardially perfused with PBS and then 4% PFA in PBS; after 18h of post-fixation in 4% PFA, brains were cryoprotected in 30% Sucrose/PBS, embedded in OCT and serially sectioned using a *Leica CM1950 cryostat* into 70 μm sections from AP +2.6 mm to −3.0 mm as previously reported (Commisso et al., 2018).

The ZsGreen reporter did not require immunodetection since the native signal was very strong. For dedicated experiments, immunostaining for SOD1 (Anti SOD1, Sigma-Aldrich, HPA001401, 1:500), misfolded SOD1 (Misfolded SOD1 (B8H10), MédiMabs, MM-0070-p, 1:250) and Fus (Anti Fus, Sigma-Aldrich, HPA008784, 1:300) was performed.

Briefly, sections were blocked in blocking solution (3% BSA, 0.3% Triton in PBS) for 2h at room temperature (rt). Primary antibody was diluted in blocking solution and incubated at 4°C for 72h. Secondary antibody (Invitrogen, Donkey anti-rabbit Alexa Fluor 568, # A10042; Donkey anti-mouse Alexa Fluor 647, # A31571; anti-guinea pig 405) was applied diluted 1:500 in blocking solution after three washing steps (PBS 20 min at rt) and incubated for 2h at rt. Subsequently sections were washed three times again and mounted using Gold Antifade Mountant (Invitrogen #P36930).

### Image acquisition

Glass slides of serially-sectioned brains were first subject to visual assessment using a Leica DMIL equipped with a 2.5x/0.07 objective; brains that were injected in the wrong location or displayed obvious macroscopic artifacts were excluded at this stage and not processed further.

The remaining set of brain sections were imaged in full using a slide-scanning microscope (Leica DMI6000B) equipped with a 5x/0.12 objective and exposure time ranging between 300 ms (405 nm) to 800ms (647 nm). All images were saved at 15304 × 28295 resolution with 16-bit depth. Images of the cortical areas immunostained for misfolded SOD1 were acquired using a Zeiss LSM710 confocal laser scanning microscope equipped with a 20x air objective. In total, 15 optical sections each 1μm-thick were acquired for each region of interest.

### Neuron mapping

A multiple software approach was devised to analyze brain section images and properly parcellate and quantify projections from cortex to LHA; the pipeline is summarized in Fig. 1A and is reported here in detail.

**Figure 1:**
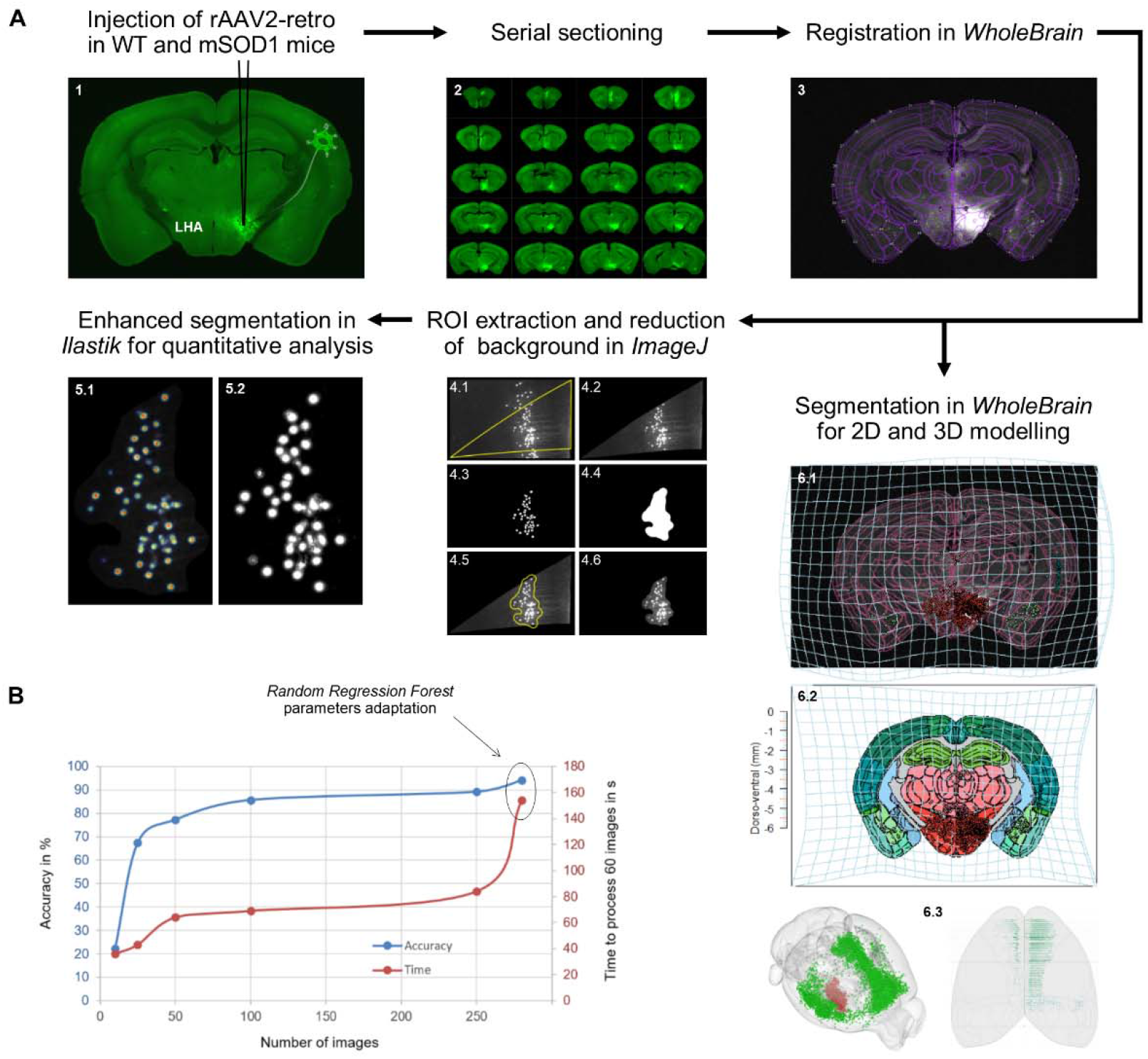
Pipeline for the quantitation of forebrain projections to LHA. A: Neurons projecting to LHA are identified by injecting rAAV-retro encoding Cre in the LHA of WT (or mSOD1) carrying a floxed ZsGreen reporter allele (panel 1). Brains are serially sectioned (2) and each section is manually registered in *WholeBrain* (3). To obtain a precise quantification of the neurons in each anatomical structure, registered sections are parcellated and background-corrected in ImageJ (panel 4) and the parcellated images are segmented using a purpose-trained *Ilastik* (density counting workflow) (5) to obtain the final neuronal counts per image. The *WholeBrain* dataset is also used to derive an independent anatomical annotation in the Allen Brain Atlas (backward warp transform field (atlas plate to section) 6.1, forward warp transformation field (section to atlas plate) 6.2) and 3D reconstructions (6.3). B: Training of *Ilastik* cell density counting. Accuracy improvement of the classifier according to the amount of images used for training. The increasing need of computing power is indicated by the time needed to process 60 Images. Accuracy was calculated by comparing randomly selected and manually counted images. Using a total of 250 images and adapting the parameters for the *Random Regression Forest*, a final accuracy of 94% was achieved.

#### Anatomical annotation

Firstly, a custom *Fiji* (Schindelin *et al.*, 2012) macro allowed extracting single brain sections from whole microscopic slide images while appending serial numbers. Hands-on registration of coronal brain sections was performed by means of *WholeBrain* package in R (Fürth *et al.*, 2018), laying its foundations on the Allen Brain mouse reference Atlas (ABA version 2011) (Lein, E.S. et al. 2007). In view of accounting for tilted sections (as compared to the original ABA coronal planes) and for biological inter-individual brain differences, a series of easily recognizable anatomical landmarks were chosen to appropriately map the regions of interest (ROIs) along the anterior-posterior (AP) axis, as shown in Supplementary Table 1. The stereotactic coordinates were determined with *openbrainmap.org*.

#### Neuron segmentation

Despite the flexibility of the parameters of the built-in segmentation functions in *WholeBrain*, a systematic underestimation of neuronal counts was noticeable in areas with high neuronal density, such as the frontal cortex, whereas a systematic overestimation of neuronal counts took place in areas with low neuronal density, due to the inclusion of segments of dendrites as purported neurons. In order to avoid the warping of the neuronal counts because of these opposing biases, a separate strategy was pursued for neuron segmentation in single brain ROIs alone. RGB images with single region outlines were generated as explained in https://gitter.im/tractatus/Lobby and imported in *Fiji* (Schindelin et al., 2012). Another *Fiji* macro was then designed to mask and resize the RGB image in order to have a perfect match of the original high-resolution image, to convert the former to HSB Stack, recover the outline of the brain ROI and use it to crop the latter.

*Ilastik* toolkit (Berg *et al.*, 2019) subsequently served a double purpose. The Pixel Classification workflow enabled to discriminate neurons (foreground) from neuron-devoid brain architecture (background); the resulting binary images (Fig. 1A, 4.3) were fed to a final *Fiji* macro to crop highly-resolved brain ROIs (Fig.1A, 4.2) to smaller ROIs containing neurons only (Fig.1A, 4.6) and exclude brain ROIs without LHA-projecting somata from further analysis. This step proved crucial to resort to a highly trained Cell Density Counting *Ilastik* suite, since brain architecture removal reduced the chance of generating false positives and drastically lessened the burden on the RAM.

### Ilastik training

For both pixel classification and cell density counting workflows, σ values of 1.0, 1.6, 3.5 and 5.0 were selected for the following features: Gaussian Smoothing, Laplacian of Gaussian, Gaussian Gradient Magnitude, Difference of Gaussian, Structure Tensor Eigenvalues, Hessian of Gaussian Eigenvalues. Twenty randomly selected whole-section images were sufficient for Pixel Classification training; batch processing generated simple-segmentation.tiff (Figure1, A, 4.3) images.

On the other hand, the Cell Density Counting classifier required a total of 250 randomly selected images for training. As displayed in figure 1B, after annotating 100 images the training effect had already reached saturation. Since accuracy of less than 90% seemed not satisfying enough, additional 150 images were annotated to tweak out a few more percentages in accuracy. Furthermore, we adapted the parameters of the implemented *Random Regression Forest* algorithm (described in more detail in Fiaschi *et al.*, 2012), namely number and maximum depth of the trees (Ntrees = 20, MaxDepth = 80), achieving a final accuracy of 94% by comparing counts to a selection of 100 randomly selected and manually counted images. Random sampling of probability maps exported upon batch image processing provided a further quality control (Fig.1A, 5.2).

### Determination of single brain region and injection site volumes

Volume measurements were performed with yet another custom macro in *Fiji*, measuring the area of individual scaled cropped.tiff images (Figure1, A, 4.2) and multiplying by the thickness of a single section (70 μm). Injection site volumes were determined by manually defining their cross-section in every brain slice at a set image contrast and once again multiplying such areas for the section thickness. Brains with injection sites bigger than 1 mm^3^, mislocalized position or displaying a virus backflow (visible in dorsal regions with respect to the LHA) bigger than 10% of the whole injection site itself were excluded from further analysis.

### Determination of misfSOD1 burden

A dedicated macro in Fiji was used to create a selection of misfSOD positive cells by applying the RenyiEntropy (Sahoo et al., 1997) thresholding and subsequent background subtraction (rolling ball radius = 20 px) and gaussian blurring (σ = 1). Afterwards, the area was measured by restoring the selection in the original image

### MRI scanning of DTI data in ALS patients

DTI scanning was performed in 72 ALS patients (58 ± 13 years, 42m, ALS-FRS-R 40 ± 6, disease duration 19 ± 15 months) and 43 controls (56 ± 9 years, 24m) according to a standardized protocol (for details see Kassubek et al., 2018). DTI data were acquired on a 3.0 T head scanner (Allegra, Siemens Medical, Erlangen, Germany). The standardized DTI scanning protocol was as follows: 49 gradient directions (b = 1000 s/mm^2^) including one b = 0 gradient directions, 52 slices, 96 × 128 voxels in-plane, slice thickness 2.2 mm, in-plane voxel size 2.2 mm x 2.2 mm, echo time 85 ms, repetition time 7600 ms. All participating patients and controls provided written informed consent for the study according to institutional guidelines. The study was approved by the Ethical Committee of the University of Ulm (reference #19/12).

### DTI analysis

The DTI analysis software Tensor Imaging and Fiber Tracking (TIFT; Müller et al., 2007A) was used for post processing and statistical analysis as previously described (Kassubek et al., 2018). In brief, stereotaxic normalization to the Montreal Neurological Institute (MNI) space was performed iteratively using study-specific templates. To map white matter microstructure, fractional anisotropy (FA) maps were calculated from the stereotaxically normalized DTI data sets of all subjects. A Gaussian filter of 8 mm full width at half maximum was applied for smoothing of FA maps for a good balance between sensitivity and specificity (Unrath et al., 2010) and FA maps were corrected for the covariate age. Tractwise fractional anisotropy statistics (TFAS) was performed by statistically comparing the FA values between the two subject groups in a given tract system (Müller et al., 2007B). The following tract systems were focussed on: a tract from orbitofrontal regions to the hypothalamus, a tract from the hypothalamus to the insula, a cingulate-hypothalamic tract, and as an ALS-reference the corticospinal tract. Consequently, a tract-of-interest (TOI) analysis allowed for quantification of microstructural alterations in these tract systems.

### Statistical analysis

The statistical analysis was performed with the GraphPad8 software suite. Comparison of the neuronal counts (absolute counts or normalized) was performed by two-way ANOVA (genotype x structure design) using Sidak’s post-hoc multiple comparisons correction; corrected p values are provided. Normalization was obtained dividing the absolute number of neurons in each structure for the total number of neurons counted in that brain and multiplying the ratio for 50,000. Comparison of total neuronal counts was performed by two-tailed unpaired Student’s t-test. Comparison of misfSOD1 burden was performed by one-way ANOVA with Tuckey’s post-hoc test.

## Results

### Enhanced semi-automated mouse brain segmentation with neuronal annotation for global quantification of projections with single-cell resolution

In order to build a quantitative and anatomically accurate map of projections from cortical and subcortical areas to the lateral hypothalamus, first we set out to establish a reliable approach to identify neurons and register their position within the structure classification of the Allen Brain Anatomical reference Atlas (Fig. 1A). The image batch for software training was obtained by injecting the retrograde rAAV2 (rAAV2-retro) into the LHA of two ZsGreen reporter mice. The latter were sacrificed 15 days later and the fixed brain was serially sectioned in 70 μm-thick sections; the whole glass slide with mounted sections was scanned using a Leica epifluorescence microscope. Each atlas plate was warped onto the brain section by defining an appropriate number of corresponding anatomical landmarks with WholeBrain (see the Methods section), thereby matching the atlas coordinates with the actual brain sections and allowing a proper anatomical annotation (Fig. 1A, 2). Single structure outlines (cortical layers, subcortical structures, e.g. fig. 1A, 4.1) were extracted using a dedicated *R* script, size-corrected and re-overlapped onto the original image in *Fiji*; the corresponding area of the original image was then cropped (Fig. 1A, 4.2), resulting in the unpacking of the original single-section into several hundreds cropped pictures (each image was coded to be later identified), resulting in >100,000 pictures per brain. In order to identify and count the number of neurons in each fragment, *Ilastik* software suite was employed. Simple foreground/background segmentations were generated by resorting to the pixel classification workflow. This allowed to automatically clear most of the brain autofluorescence background in *Fiji* (Fig. 1A, 4.6). This step enabled the improvement of the discrimination of few ZsGreen+ neurons in ROI containing many more dendrite stretches and artifacts, and also reduced the computational burden placed on *Ilastik cell density* classifier and therefore minimizing the chance of generating artifactual counts. Ilastik cell density classifier was trained with 250 randomly selected ROI images that were annotated by a human operator; the accuracy of software reached 86% after 100 images and achieved a final 94% peak (Fig. 1B) by improving the performance of the *Random Regression Forest* algorithm with 150 additional images and tuning its parameters (trees number and depth; see Methods). The whole procedure was then sequentially implemented so that neuronal counts for each picture from every section were annotated and logged together with the anatomical location within the hemisphere and the brain section as well as with the rostro-caudal serial number of the section itself.

### Disrupted cortico-hypothalamic projections in symptomatic mSOD1 ALS mice

We exploited our enhanced registration-segmentation-annotation pipeline to establish whether the large-scale architecture of the projections to LHA was altered in the mSOD1 mice at a symptomatic stage (i.e., when body weight loss has appeared and clinical score is 2). To this aim, 20 mSOD1/ZsGreen+ and 20 WT/ZsGreen+ mice were injected at the age of P95 with rAAV2-retro (Tervo et al., 2016) encoding for Cre under the human synapsin promoter in the Lateral hypothalamic area (LHA; Fig. 2A). All neurons projecting to the site of injection would take up the rAAV2-retro, express the Cre recombinase and appear ZsGreen+. Mice were sacrificed 15 days later. Upon serial sectioning, a series of quality control criteria were applied and brains displaying i) improper location of the injection site ii) injection volume larger than the boundaries of the LHA iii) large number of neurons infected along the thread of the injection capillary iv) low (<25,000) overall number of ZsGreen+ neurons were excluded from further analysis. Six brains for each genotype were then considered for further quantification. These brains were randomly divided in two cohorts: cohort 1 was subjected to *WholeBrain* parcellation and cohort 2 was left for confirmation (see below). Once cohort 1 brains were processed through the *WholeBrain*-registration and annotation pipeline, we identified an average of about 50,000 neurons projecting to the LHA in WT mice; when the brainstem was excluded (because anatomical annotation of brainstem nuclei was judged unreliable based on visual inspection of the *WholeBrain*-overlapped images), we identified 84 cerebral areas projecting to the LHA (full list with absolute and normalized neuronal counts is provided in Supplementary Table 2). Projections were identified from a large fraction of the forebrain, with the largest component provided by limbic areas such as Anterior Cingulate Area (ACA), Prelimbic Cortex (PL), InfraLimbic Area (ILA), Agranular Insula (AI), as well as from subcortical structures belonging to the limbic system such as Basolateral and Basomedial Amygdala (BLA and BMA, respectively). Intriguingly, substantial projections were identified from motor areas (primary and secondary), from primary sensory areas (gustatory, auditory, olfactory/piriform, visual cortex) and from basal ganglia (Dorsal Putamen, DP), underscoring the breath of integration taking place in LHA. The same structures appeared to project to LHA from ipsilateral and contralateral hemispheres, although the contralateral contribution appeared to be substantially smaller (approx. 40,000 neurons from the ipsilateral hemisphere vs 10,000 from the contralateral in WT animals).

**Figure 2:**
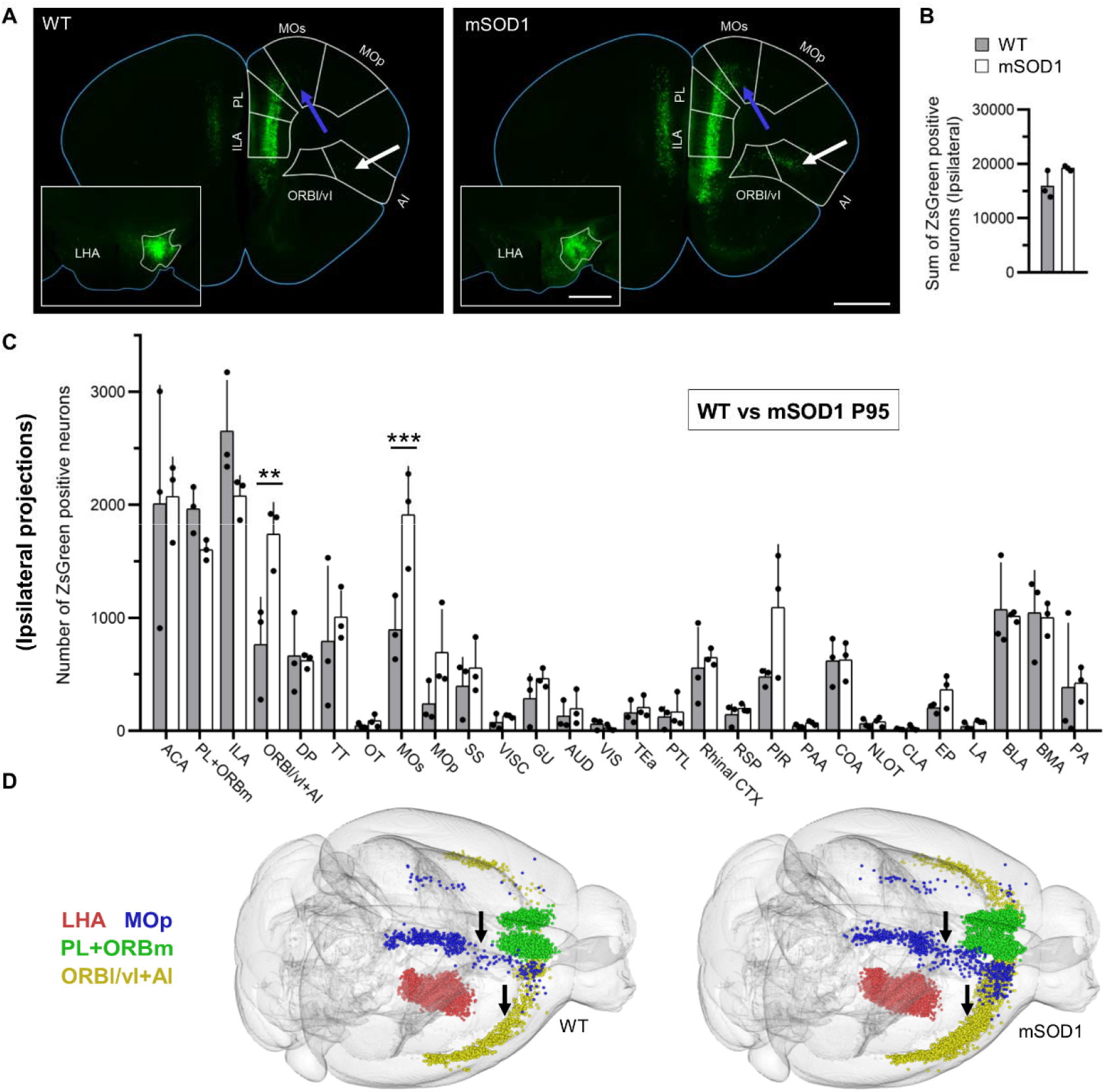
Altered cortico-hypothalamic projection pattern in mSOD1 mice at P95. A: Representative frontal brain sections of WT and mSOD1 mice depicting projections from ORBl/vl+AI (significantly increased, arrow), PL+ORBm, ILA, ACA, MOs and MOp to LHA. Inset: representative injection sites in LHA for WT and mSOD1 mice. White outlines represent LHA boundaries. B: Sum of neurons projecting from selected 28 areas. No difference is detected in the summarized number of neurons projecting to LHA in WT and mSOD1 from the ipsilateral hemisphere (n=3). C: Quantification of the number of neurons (normalized for total neuronal counts) projecting to LHA from 28 brain structures in WT and mSOD1. A significant increase in projections from ORBl/vl+AI (p=0.0012) and from MOs (p=0.0006) is detected. D: Representative *WholeBrain* volume reconstructions of neurons projecting to LHA in WT and mSOD1 mice. Expansion of projections from MOs (blue) and ORBl/vl+AI (yellow) are visible (arrows). Bars show mean ± SD. Scalebars 1 mm. \**p* < 0.05, \*\*\**p* < 0.001, \*\*\*\**p*< 0.0001.

In order to obtain a pertinent comparison (i.e., excluding areas with counts <100 neurons) of projections in WT and mSOD1 mice, we contrasted the absolute neuronal counts in 28 areas (accounting for approx. 50% of the total projections, maintaining ipsi and contralateral dataset distinct; PL and orbitomedial cortex as well as lateral/ventrolateral orbitofrontal cortex -ORBl/vl- and AI were grouped together due to the uncertainty in establishing their borders and in agreement with functional similarity. While the overall pattern of projections to LHA in symptomatic mSOD1 mice was comparable to the WT ones, the cumulative absolute counts of ZsGreen+ neurons for the 28 structures were significantly increasedin the mSOD cohort considering the absolute counts (p=0.02; SI Appendix, Fig. 1A). We contrasted the absolute counts for each of the 28 areas in WT and mSOD1 mice (SI Appendix, Fig. 1B). Two-way ANOVA revealed a significant effect of genotype (F_1,112_=105.70, p<0.0001) and post-hoc analysis (Sidak’s multiple comparisons test) revealed significant expansion of the projections from ACA (p<0.0001), ILA (p=0.009), ORBl/vl+AI (p<0.0001), Tenia Tecta (TT; p=0.0365), secondary motor cortex (MOs, p<0.0001) and piriform cortex (PIR; p=0.0005). In order to compensate for the variable total number of infected neurons in each individual brain, we then normalized the absolute counts of neurons projecting to LHA to a nominal of 50,000 neurons, so to compute the relative contribution of each area to the total input to LHA. Upon normalization, the 28 areas in object amounted for a comparable total number of neurons (Fig. 2B). Two-way ANOVA on normalized data revealed once again a significant effect of genotype (F_1,112_= 7.31, p=0.0079) and post-hoc analysis revealed a significant increase in projections from ORBl/vl+AI (p=0.0012) and from MOs (p=0.0006; Fig. 2C; also visualized in the 2D projections, SI Appendix, Fig. 1C-D as well as in the 3D Wholebrain reconstruction, Fig. 2D). The remaining 26 structures displayed only non-significant trends toward expanded projections (PIR: p=0.21; ILA: p=0.32) or no difference in the size of the neuronal population projection to LHA (Fig. 2C). When the projections from the contralateral hemisphere were contrasted, we detected an effect of genotype in the absolute counts (two-way ANOVA F_1,112_= 31.49; p<0.0001) that was traced in post-hoc test (Sydak’s) to a significant expansion of projections from ACA (p=0.0008), ILA and PL+ORBm (both p<0.0001) (SI Appendix, Fig. 2A-B). However, when the absolute counts were normalized to account for the total number of ZsGreen^+^ neurons in each brain, no significant differences were identified between the genotypes (two-way ANOVA, F_1,112_= 0.31, p=0.57; *SI Appendix*, Fig. 2C).

Since mice with neurodegenerative phenotypes may display cortical atrophy (Petrik et al., 2007), we used the *WholeBrain* registration to measure the volume of each cortical structure and compensate for eventual volume loss. Indeed, we identified a significant loss of volume in mSOD1 mice (two-way ANOVA F_1,112_=28.80, p<0.0001) which was traced by post-hoc test (Sidak’s.) to the atrophy of MOp (p=0.008) and SS (p<0.0001) with only a trend seen for MOs (p=0.33) (SI Appendix, Fig. 3). Notably, ORBl/vl+AI and MOs did not display significant changes in volume.

Thus, in symptomatic mSOD1 mice (displaying the hypermetabolic phenotype), a substantial remodeling of cortico-hypothalamic projections takes place, in particular from ORBl/vl+AI and MOs.

### Validation of cortico-hypothalamic projections abnormalities in independent operator-registered brain cohorts

Since the identification of different cortical and subcortical areas was based on the assumption of the overall similarity of the brain of mSOD1 and WT mice, we took into consideration the possibility that biases in the identification of anatomical areas could have been introduced by *WholeBrain* (e.g., because of atrophy), thereby mis-attributing neurons to cortical areas. Therefore, we set out to validate our findings using the brain cohort 2 (n=3 for WT and mSOD1; LHA injection; SI Appendix, Fig. 4A). These brain images were subjected to manual registration, i.e. each anatomical structure and cortical area was identified by a genotype-blind operator using structural landmarks (SI Table 1) and individually cropped out of each image of each section of the brain. This approach was substantially slower than the registration by *WholeBrain* (it required >500h of operator’s work time for the whole dataset to be registered and cropped). The individual pictures cropped out of the brain images were then subject to neuronal counts by *Ilastik* (since the accuracy and reproducibility of this step was already established and quantified). In the manually-annotated dataset, the total number of neurons projecting to LHA was found to be not significantly different in the mSOD1 compared to the WT (SI Appendix, Fig. 4B). First, we considered the absolute neuronal countings: the contrast of the two genotypes (ipsilateral hemisphere) revealed a statistically significant difference (two-way ANOVA F_1,112_=6.28; p=0.0136) which was further explored in post-hoc comparison (Sydak’s) revealing a significant expansion of projections from ORBl/vl+AI in mSOD1 mice (p=0.035) (SI Appendix, Fig. 4C). Upon normalization we once again detected a significant effect of genotype (two-way ANOVA F_1,112_=23.78; p<0.0001) with post-hoc revealing significant differences in ORBl/vl+AI (p<0.0001), ILA (p=0.001) and PIR (p<0.0001) (SI Appendix, Fig. 4D). When the projections from the contralateral hemisphere were considered, no overall, genotype specific or area correlated effect was found in the absolute counts (SI Appendix, Fig.5A-B); however, upon normalization, a statistically significant loss of projections from PL+ORBm was identified (p<0.0001), together with a strong trend toward increased projections from ORBl/vl+AI (p=0.072) (SI Appendix, Fig. 5C). Thus, the manually-registered dataset displayed substantial similarities with the *WholeBrain*-registered dataset (expansion of the projections from ORBl/vl+AI) but also a major discrepancy: increased projection from ILA to LHA in the manual dataset and no difference in MOs projections, whereas the opposite was detected by *WholeBrain*. Upon closer inspection of MOs projection to LHA we realized that the boundaries of MOs and ILA display a high degree of subjectivity when manually drawn and it is highly likely that MOs neurons were allocated to adjacent areas in the same way ILA was moved to neighboring areas in the dataset parcellated by the human operator.

Nevertheless, the manually-registered independent dataset confirmed the expansion of projections from AI/ORBvl to LHA.

### Unaltered projections to LHA in early presymptomatic mSOD mice

Remodelling of large-scale cortical circuits has been previously reported (Commisso et al., 2018) and appears to progress over time; nevertheless, early signs of increased projections from somatosensory cortex (SS) to primary motor cortex (MOp) were detected already at early presymptomatic stages. We set out to verify if the increased projections from AI/ORBvl appeared only late, in coincidence with the appearance of the metabolic phenotype, or if they were present already at presymptomatic stage (and possibly of developmental origin). To this aim, we injected rAAV2-retro in the LHA of WT or mSOD1 at the age of P25 (SI Appendix, Fig. 6A) and sacrificed the mice at P40. After quality control, we assessed the forebrain projections to LHA in the two genotypes (n=3 for each). The overall pattern of projections to LHA at P25 was comparable to the one observed at P95 and there was no difference in the absolute number of neurons projecting to LHA in the two genotypes (SI Appendix, Fig. 6B). The direct contrast of the two genotypes for the absolute and relative contribution of 28 selected ipsilateral structures revealed a genotype effect in absolute (two-way ANOVA: F_1,112_= 12.55; p=0.006) but not in the normalized dataset (two-way ANOVA: F_1,112_= 1.83; p=0.1779). However, in both cases the post-hoc comparison did not reveal any significant difference between the two genotypes in any of the considered structures (SI Appendix, Fig. 6C-D). Likewise, no statistical difference was found in the sum of projection from the contralateral hemisphere in absolute counts (SI Appendix, Fig. 7A), but significant increase in projections from ACA was identified (p=0.0078) (SI Appendix, Fig. 7B). Nevertheless, this difference disappeared after normalization (SI Appendix, Fig. 7C) and no differences are perceptible in the 3D brain reconstructions (SI Appendix, Fig. 7D). These findings suggest that altered AI/ORBvl connectivity to LHA arises during disease progression and it is not a developmental feature of mSOD1 brain architecture.

### ALS-related pathology in the AI/ORBvl of mSOD1 mice

We further investigated the degree of involvement of cortico-hypothalamic projections in ALS by assessing the burden of misfolded SOD1 pathology in cortical areas projecting to LHA, in particular those displaying altered connectivity compared to WT animals. We injected WT or mSOD1 mice with rAAV2-retro in LHA at P95 and sacrificed them at P110. Brain sections encompassing MOp, AI/ORBvl, PL+ORBm, (SS) were immunostained with an antibody recognizing the misfolded conformation of SOD1 (B8H10, Pickles et al., 2013) (Fig. 3A-B). The total burden of misfSOD1 was different across the cortical areas (one-way ANOVA F_3,20_=51.26, p<0.0001; MOp vs. ORBl/vl+AI p<0.0001, MOp vs. PL+ORBm p<0.0001, SS vs. ORBl/vl+AI p=0.0014, ORBl/AI vs. PL+ORBm p=0.0006), with MOp displaying the highest burden compared to SS but with ORBl/vl+AI displaying a significantly higher burden than areas showing no increase in projections such as SS or PL+ORBm (Fig. 3A-B). Interestingly, neurons projecting from MOs to LHA display high levels of misfSOD1 accumulation, whereas this is not true for neurons projecting from PL+ORBm and ORBl/vl+AI (Fig.3A).

**Figure 3:**
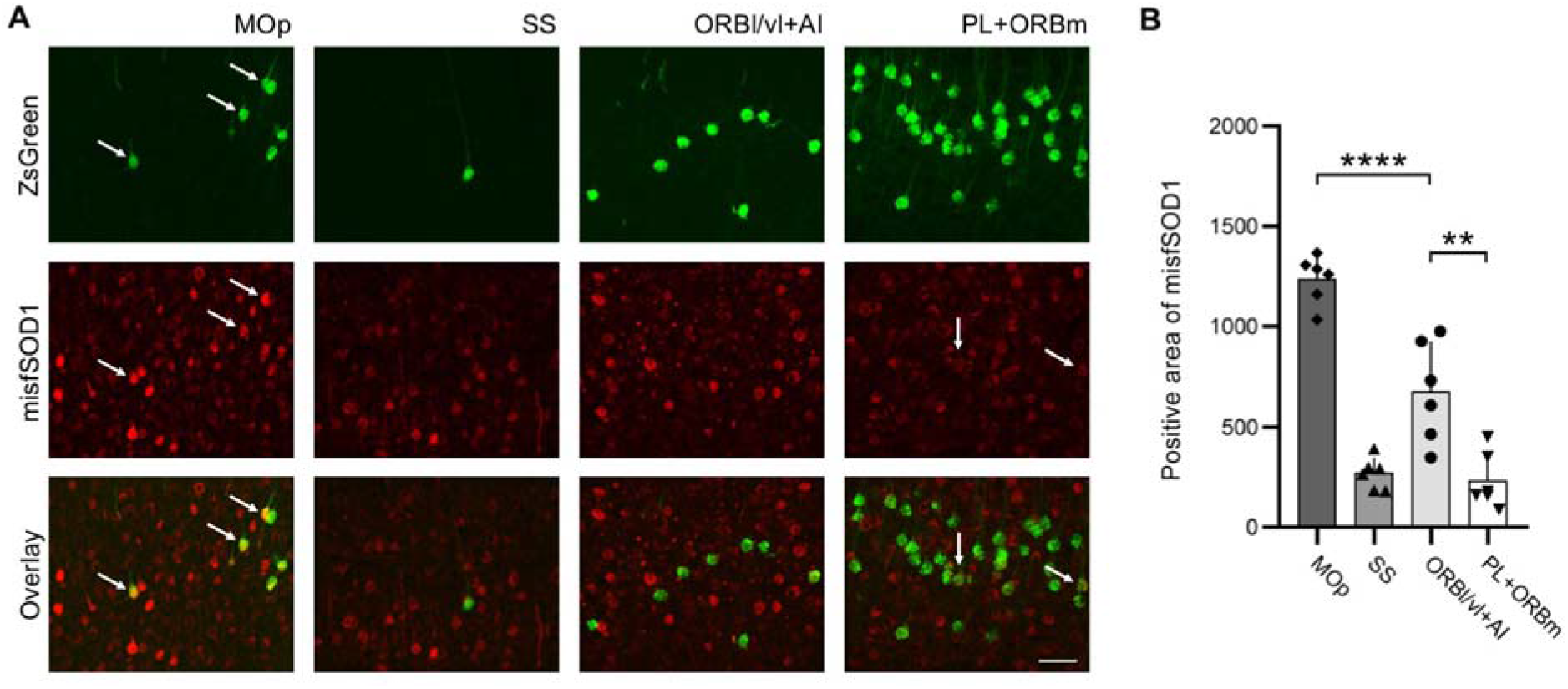
Significant misfolded SOD1 burden in ORBl/vl+AI in mSOD1 mice. A: Representative images of misfSOD1+ cells in MOp, SS, ORBl/vl+AI and PL+ORBm; ZsGreen^+^ neurons projecting to LHA show a strong misfSOD1 burden in MOp (arrows). Other structures show no double positive neurons (SS,ORBl/vl+AI) or weak mSOD burden (PL+ORBm; arrows). B: Burden of misfolded SOD1 (detected by the B8H10 antibody) in MOp, SS, ORBl/vl+AI and PL+ORBm. ORBl/vl+AI displays a burden significantly higher than SS or PL (p=0.0014 and p=0.0006 respectively; areas with unchanged connectivity to LHA) but significantly lower than MOp (p<0.0001). Bars show mean ± SD. Scalebars 50 μm. \*\**p* < 0.01, \*\*\*\**p*< 0.0001.

### Distinct pattern of altered cortical projections to LHA in Fus mice

We further explored the alteration of projections to LHA in ALS in an independent ALS model, the kiFUS mouse. Of note, the kiFUS mouse differs from the mSOD1 mouse because it does not display an overt body-weight phenotype (Scekic-Zahirovic et al., 2017) and shows a more limited loss of spinal motoneurons and slower progression (Ouali-Alami et al., 2020). We proceeded with a similar experimental design as for the mSOD1 mice; however, since kiFUS mice display spinal motoneuron loss not before 9-10 months of age, we considered the P255 as “early symptomatic” stage. Therefore, kiFUS mice and their own WT littermates were injected at P255 into the LHA (Fig. 4A and sacrificed at P270 (n=4 for each group). Once again, there was no difference in the overall number of neurons projecting to LHA across the forebrain in absolute counts (SI Appendix, Fig. 8A). When the absolute neuronal counts of each of the 28 areas were contrasted, we detected a significant effect of the genotype (two-way ANOVA F_1,166_=8.583; p=0.0039) and post-hoc comparison revealed a significant decrease in kiFUS in projections from ACA (p=0.0385), MOs (p<0.0001) and strong trends for MOp (p=0.1543) and SS (p=0.0848) (SI Appendix, Fig. 8B). Upon normalization for 50,000 neurons, a significant difference in MOs was still detected (p<0.0001) together with strong trends for ACA (p=0.1639) and MOp (p=0.136) (Fig. 4B-C and 2D models in SI Appendix, Fig. 8C-D). Interestingly, a significant genotype effect was also found in the contralateral hemisphere both in absolute (SI Appendix, Fig. 9A-B) and normalized data (SI Appendix, Fig. 9C) (two-way ANOVA, F_1,166_= 6.57; p=0.0112 and F_1,166_= 4.48 p=0.0356, respectively) and differences were revealed by post-hoc analysis in loss of projections from ACA (p=0.0121 and p=0.0137 in absolute counts and normalized, respectively) and ILA (p=0.0474) in the absolute counts only. Of note, projections from AI/ORBvl were not affected (neither from ipsilateral nor from contralateral). The significant loss of MOs neurons projecting to LHA was especially affecting the most rostral part of MOs, as demonstrated by the 3D reconstruction model of the mouse brain (Fig. 4D). Thus, distinct ALS models with divergent metabolic phenotype display different patterns of altered projections to LHA.

**Figure 4:**
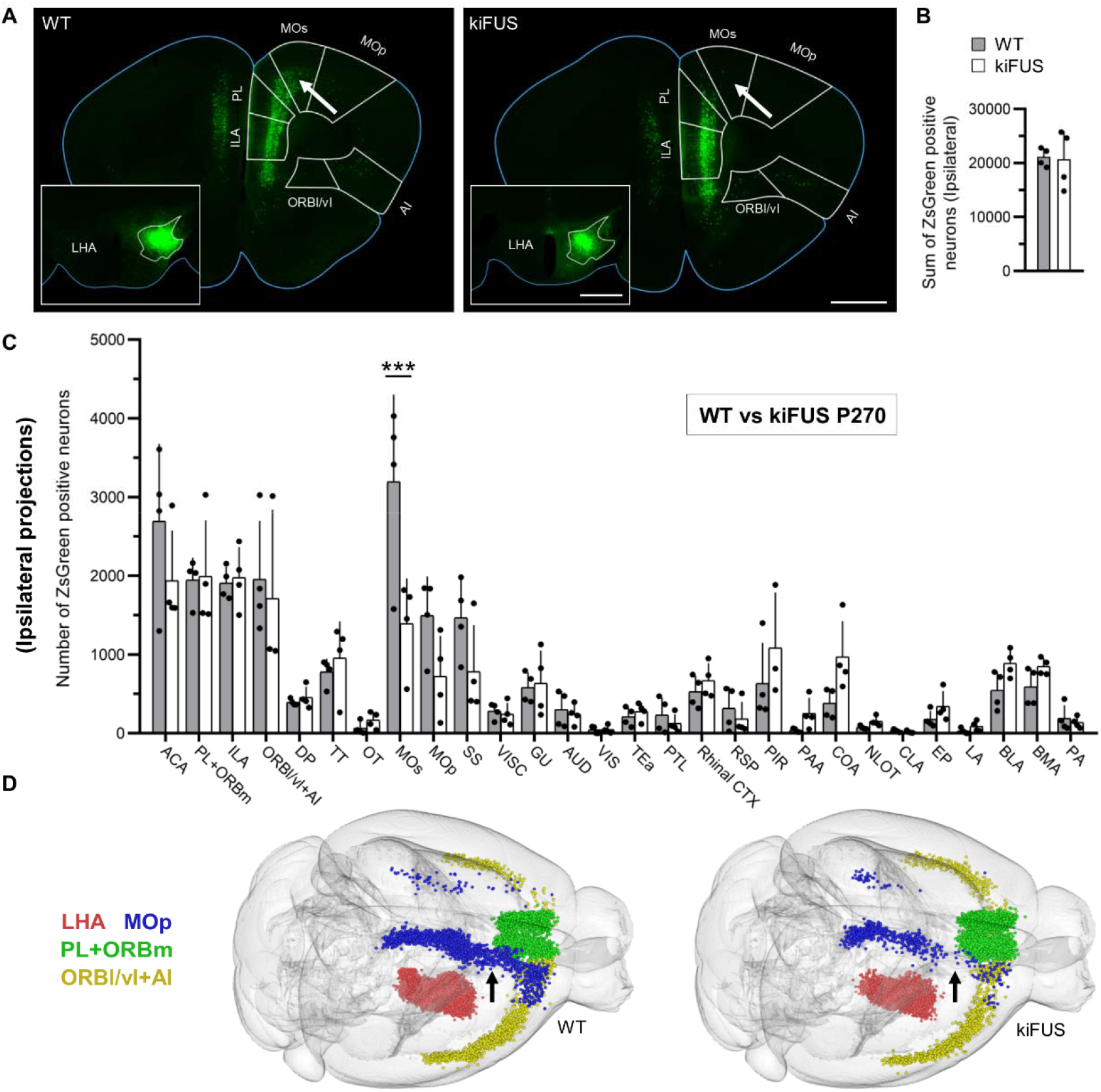
Altered cortico-hypothalamic projection pattern in kiFUS mice at P270. A: Representative frontal brain sections of WT and kiFUS mice depicting projections from MOs (significantly decreased, arrow), PL+ORBm, ILA, ACA, and MOp to LHA. Inset: representative injection sites in LHA for WT and kiFUS mice. White outlines represent LHA boundaries. B: Sum of neurons projecting from selected 28 areas. No difference is detected in the summarized number of neurons projecting to LHA in WT and kiFUS from the ipsilateral hemisphere (n=4). C: Quantification of the number of neurons (normalized for total neuronal counts) projecting to LHA from 28 brain structures in WT and kiFUS. A significant decrease in projections from MOs (p<0.0001) and trend toward decrease projection from ACA (p=0.1639) and MOp (p=0.136) is detected (n=4). D: Representative *WholeBrain* volume reconstructions of neurons projecting to LHA in WT and kiFUS mice. Loss of projections from the anterior part of MOs (blue) is visible. Bars show mean ± SD. Scalebars 1 mm. \**p* < 0.05, \*\**p* < 0.01, \*\*\*\**p* < 0.001.

### DTI-MRI reveals altered orbitofrontal-hypothalamic tract in ALS patients

Finally, we set out to investigate if the selective disturbance in cortico-hypothalamic projections from AI, ORBvl and PL observed in the mSOD1 mice could also be observed in human ALS patients. To this aim, we elected to use a 3T DTI-MRI dataset involving 83 ALS patients and 64 healthy subjects (Kassubek et al., 2018). Four white-matter tracts were taken into consideration: the orbitofrontal-hypothalamic tract, the insular-hypothalamic tract, the cingulate-hypothalamic tract (Fig. 5) and as reference the corticospinal tract which is known to be substantially altered in ALS patients (Kassubek et al., 2018); the three tracts converging on the hypothalamus were selected because they provided the closest possible match (when accounting for the different anatomy) for the structures investigated in the mouse model and because of their reproducible and unequivocal identification in DTI datasets. Whole brain FA showed a slight reduction in ALS patients. A significant FA reduction (p < 0.05) could be detected in the tract from orbitofrontal regions to the hypothalamus. No significant FA alteration could be detected in a tract from the hypothalamus to the insula and a cingulate-hypothalamic tract. In the ALS-reference tract (the corticospinal tract) significant FA reduction (p < 0.01) could be observed, as previously detected (Kassubek et al., 2014; Kassubek et al., 2018), in a cerebellar reference (where no ALS affection can be anticipated), no significant FA alteration was detected. Thus, disturbances of cortical projections to hypothalamus are not unique characteristics of the mSOD1 murine model of ALS but constitute a previously unrecognized architectural phenotype shared by human ALS patients.

**Figure 5:**
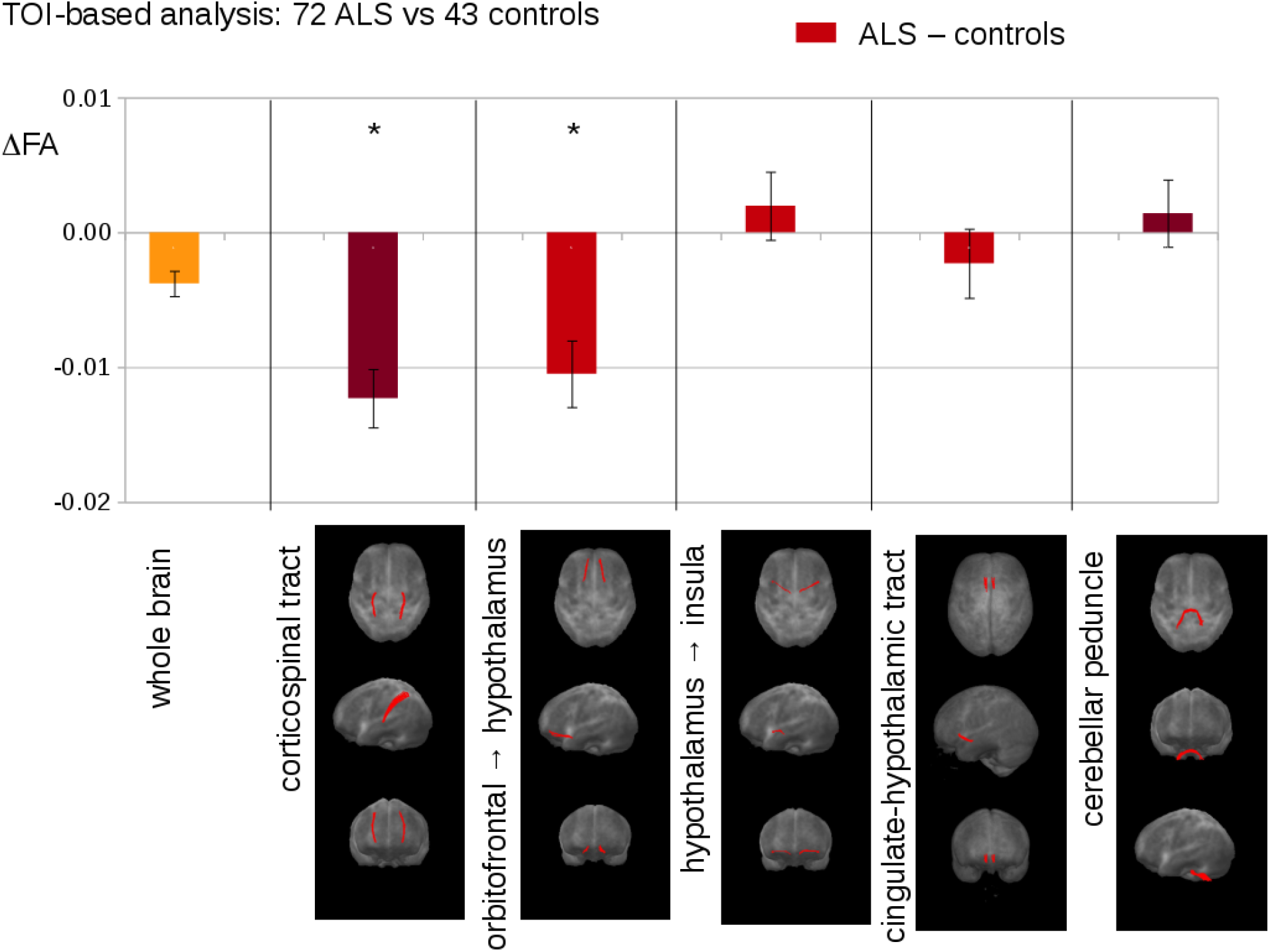
Tract of Interest (TOI)-based analysis of DTI data from 83 ALS patients vs 64 controls. Upper panel: differences of mean fractional anisotropy (FA) values between ALS patients and controls. Lower panel: projectional views (axial, coronal, sagittal) of tract systems used for TOI analysis. TOI – tract-of-interest; **p < 0.05; *p < 0.001; error bars are the standard error of the mean (SEM).

## Discussion

Here we provide converging evidence from ALS mouse models and human imaging datasets of the involvement of large-scale projections to LHA in ALS, in particular the disruption of connections from Orbitofrontal Cortex and/or Agranular Insula to LHA. This effect is independent of systematic biases introduced by software or human operators in the registration, does not appear to be an intrinsic developmental defect and it is not detected in an independent ALS model devoid of metabolic phenotypes. The findings in the murine model display striking similarity to the disruption of the orbitofrontal-hypothalamic tract identified in a large dataset of DTI-MRI of ALS patients.

Hypermetabolism in ALS has been repeatedly reported in human patients (Steyn et al., 2018; Jesus et al., 2018; Ahmed et al., 2016; Desport et al., 2005; Bouteloup et al., 2009) as well as in some, but not all, ALS murine models (Dupuis et al., 2004) and, notably, it appears to develop over time and may predate the clinical onset of disease by several years (Peter et al., 2017; Mariosa et al., 2017). Nevertheless, the mechanisms for this clinical manifestation of ALS are unclear.

Direct involvement of the hypothalamus has been hypothesized on the basis of the role of this structure as the main central controller of energy homeostasis, feeding and satiety (Vercruysse et al., 2018), has been supported by evidence of reduced hypothalamic volume in ALS patients (Gorges et al., 2017) and by the detection of ALS-related TDP-43 pathology in a subset of ALS patients hypothalami (Cykowski et al., 2014). Neurochemical abnormalities, such as reduced MCH expression, have been reported in mutant SOD1 murine models (Vercruysse et al., 2016). However, these signs of intrinsic hypothalamic disturbance are not mutually exclusive with the hypothesis that large-scale hypothalami circuits may be dysfunctional, too. Disruption within the larger circuit regulating hypothalamic function, which include insular, motor, orbitofrontal and limbic areas, may contribute to the hypothalamic dysregulation by altering the equilibrium of the different inputs. Indeed, changes in neuronal activity in hypothalamus may produce substantial modifications of the neurochemical identity of local subpopulation, including switch in type of neurotransmitter released (Meng et al., 2018). Here we demonstrate that besides any pathology intrinsic to the hypothalamus, the disease pathophysiology involves a substantial quantitative change in the inputs that hypothalamus receives, particularly from agranular insula and ventrolateral orbitofrontal cortex. This remodeling process coincides with the onset of body weight loss in mSOD1.

Notably, the circuit involving projections from orbitofrontal cortex and agranular insula to lateral hypothalamus is conserved from mice and rats (Floyd et al., 2001; Hoover et al., 2011) to marmosets (Roberts et al., 2007) and macaques (Ongür et al., 1998) to humans, thus validating mouse as a model organism. Nevertheless, the murine orbitofrontal cortex (including ventral and ventrolateral) and agranular insula are considered homologous of orbitomedial cortex in primates (Price, 2007), so that the study the orbitofrontal-LHA tract in our DTI tracing study is located more medially than the ORBl/vl+AI complex in the mouse brain.

Remarkably, altered projections from AI/ORBvl in mSOD1 mice correspond to reduced FA in the orbitofrontal-hypothalamic (but not insular-hypothalamic) tract in humans. Although it is noteworthy that abnormalities appear in the same brain structures in animal models and patients with ALS, the closer examination of the findings reveal that in the murine model an actual expansion of projections is detected whereas decreased FA in DTI is usually interpreted as loss of integrity of axons or myelinated tracts (Alexander et al., 2007). However, it must be stressed that SOD1^(G93A)^ displays a fast progression and becomes symptomatic at a comparatively young age, when the potential for compensatory sprouting after degeneration may be larger than in older human patients (hence, the expansion of the projections). Thus, in human patients the DTI may reveal a change in microstructural integrity resulting from axonal loss but would be unable to detect any increase in the size of axonal arborization.

To the best of our knowledge, there is no demonstration that the orbitofrontal cortex and/or AI can directly regulate metabolic rates. It must be emphasized, though, that lesion studies of these structures have not included a rigorous, long-term testing of body-weight dynamics and inactivation studies have been performed only in short-term settings (Kobayashi, 2011; Izquiredo, 2017; Rolls, 2004). On the other hand, OFC and AI are involved in processing food characteristics, such as taste, smell and texture (Seabrook et al., 2020) and display a differential response to caloric content of food depending on the satiety of the subject (at least in humans: Schur et al., 2009; Suzuki et al., 2017). Thus, it is conceivable that altered connectivity between vlORB/AI and LHA may provide the latter with incorrect inputs regarding the nutritional content of food, contributing to the dysregulation of body metabolic rates. At least in human subjects, reduced volume of grey matter and altered microstructural integrity (measured by apparent diffusion coefficient) in orbitofrontal cortex have been observed in obese patients (Raji et al., 2010; Walther et al., 2010; Alkan et al., 2008), strengthening the link between dysfunction in orbitofrontal cortex and body weight.

Unexpectedly, our retrograde projection mapping strategy has consistently identified projections from primary and secondary motor cortex to LHA, which appear to be affected by the disease process both in mSOD1 and in kiFUS. This connectivity appears to be reciprocal, since LHA projections to motor cortices have also been reported (Commisso et al., 2018). These connections cannot be easily discounted as artifacts due to the proximity of LHA to the corticospinal tract, since they appear to originate from both ipsi and contralateral hemispheres. It is conceivable that LHA may receive inputs relaying the average motor activity performed or planned and may adjust energy balance accordingly; disruption of connectivity between motor cortex and hypothalamus may leave the latter “free running”, without a proper estimate of current motor activity. Alternatively, it is possible that any pathogenic event in motor cortex (propagating prionoids or abnormal activity patterns) may propagate to hypothalamus and drive its dysfunction and degeneration (as suggested for spinal cord; Burg et al., 2020).

Taken together, our findings suggest that disruption of large-scale circuits providing input to LHA may contribute to generate the metabolic phenotype observed in ALS; in particular, we have identified the orbitofrontal-hypothalamic tract as a site of convergence of mouse projection data and human tracing data, which may open up an independent approach to evaluate non-motor prognostic features in ALS.

## Acknowledgement

FR is supported by the Thierry Latran Foundation (projects “Trials” and “Hypothals”), by the Radala Foundation, by the Deutsche Forschungsgemeinschaft (DFG, German Research Foundation)- Project-ID 251293561 – Collaborative Research Center (CRC) 1149 and with the individual grant no. 431995586 (RO-5004/8-1) and no. 443642953 (RO5004/9-1), by the Cellular and Molecular Mechanisms in Aging (CEMMA) Research Training Group and by BMBF (FKZ 01EW1705A, as member of the ERANET-NEURON consortium “MICRONET”). We thank the colleagues of the German Center for Neurodegenerative Diseases (DZNE) for helpful comments. SA and DB are members of the International Graduate School in Molecular Medicine at Ulm University; DB is part of the Graduate School in Cellular and Molecular Mechanisms in Aging at Ulm University.

## Supplementary Information

***Supplementary Table 1***: Anatomical landmarks used for mapping the brain section from anterior to posterior. The following structures provided distinctive optical characteristics for mapping: AOB = Accessory olfactory bulb; DP = Dorsal peduncular area; fa = corpus callosum, anterior forceps; ccg = genu of corpus callosum; aco = anterior commissure, olfactory limb; act = anterior commissure, temporal limb; SCH = Suprachiasmatic nucleus; NLOT = Nucleus of the lateral olfactory tract; ME = Median eminence; opt = optic tract; int = internal capsule; DG = Dentate gyrus; CA3so = Field CA3, stratum oriens; MM = Medial mammillary nucleus.

***Supplementary Table 2***: List of structures of the brain projecting to LHA. For each structure, the absolute number of neurons projecting to LHA is reported as mean ± SD and independently for ipsi- and contralateral hemispheres.

***Supplementary Fig.1***: Altered cortico-hypothalamic projection pattern in mSOD1 mice at P95, absolute counts ipsilateral. A: Sum of neurons projecting from selected 28 areas. Significantly increased projections from ipsilateral hemisphere in mSOD mice to LHA (p=0.02; n=3). B: Quantification of the absolute number of neurons projecting to LHA from 28 brain structures in WT and mSOD1. A significant increase in projections from ACA (p<0.0001), ILA (p=0.009), ORBl/vl+AI (p<0.0001), Tenia Tecta (TT; p=0.0365), secondary Motor Cortex (MOs, p<0.0001) and Piriform cortex (PIR; p=0.0005). C: Representative *WholeBrain* ipsilateral side view reconstructions of neurons projecting to LHA in WT and mSOD1 mice. Expansion of projections from MOs are visible (arrows). D: Representative *WholeBrain* cortical top view reconstructions of neurons projecting to LHA in WT and mSOD1 mice. Expansion of projections from MOs are visible (arrows). Bars show mean ± SD. *p < 0.05, **p < 0.01, ***p < 0.001, ****p< 0.0001.

***Supplementary Fig.2***: Altered cortico-hypothalamic projection pattern in mSOD1 mice at P95, contralateral hemisphere. A: Sum of neurons projecting from selected 28 areas. No difference is detected in the summarized number of neurons projecting to LHA in WT and mSOD1 from the contralateral hemisphere (n=3). B: Quantification of the absolute number of neurons projecting to LHA from 28 brain structures in WT and mSOD1. A significant increase in projections from ACA (p=0.0008), PL+ORBm and ILA (both p<0.0001) is detected. C: No difference is detected in the quantification of the normalized number of neurons projecting to LHA from 28 brain structures in WT and mSOD1. Bars show mean ± SD. \**p* < 0.05, \*\**p* < 0.01, \*\*\**p* < 0.001, \*\*\*\**p*< 0.0001.

***Supplementary Fig.3***: Primary motor cortex and somatosensory cortex atrophy in P95 mSOD mice. Volumetric comparison of 28 areas in WT and mSOD. Significant atrophy was detected in MOp and SS. Bars show mean ± SD. \**p* < 0.05, \*\**p* < 0.01, \*\*\**p* < 0.001, \*\*\*\**p*< 0.0001.

***Supplementary Fig.4***: Altered cortico-hypothalamic projection pattern in manual registered mSOD1 mice at P95, ipsilateral. A: Left: representative injection sites in LHA for WT and mSOD1 mice. White outlines represent LHA boundaries. Middle and right: Representative frontal brain sections of WT and mSOD1 mice depicting projections from ORBl/vl+AI (significantly increased, arrow), PL+ORBm, ILA, ACA, MOs and MOp to LHA. B: Sum of neurons projecting from selected 28 areas. No difference is detected in the summarized number of neurons projecting to LHA in WT and mSOD1 from the ipsilateral hemisphere (n=3). C: Quantification of the absolute number of neurons projecting to LHA from 28 brain structures in WT and mSOD1. A significant increase in projections from ORBl/vl+AI (p=0.035) is detected. D: Quantification of the normalized number of neurons projecting to LHA from 28 brain structures in WT and mSOD1. A significant increase in projections from ILA (p=0.001), ORBl/vl+AI (p<0.0001) and PIR (p<0.0001) is detected. Bars show mean ± SD. Scalebars 1 mm. \**p* < 0.05, \*\**p* < 0.01, \*\*\**p* < 0.001, \*\*\*\**p*< 0.0001.

***Supplementary Fig.5***: Altered projection pattern in manual registered mSOD1 mice at P95, contralateral. A: Sum of neurons projecting from selected 28 areas. No difference is detected in the summarized number of neurons projecting to LHA in WT and mSOD1 from the contralateral hemisphere (n=3). B: No difference is detected in the quantification of the absolute number of neurons projecting to LHA from 28 brain structures in WT and mSOD1. C: Quantification of the normalized number of neurons projecting to LHA from 28 brain structures in WT and mSOD1. A significant increase in projections from PL+ORBm (p<0.0001) is detected. Bars show mean ± SD. \**p* < 0.05, \*\**p* < 0.01, \*\*\**p* < 0.001, \*\*\*\**p*< 0.0001.

***Supplementary Fig.6***: Unaltered cortico-hypothalamic projection pattern in mSOD1 mice at P25, ipsilateral. A: Left: representative injection sites in LHA for WT and mSOD1 mice. White outlines represent LHA boundaries. Middle and right: Representative frontal brain sections of WT and mSOD1 mice depicting a similar pattern of neurons. B: Sum of neurons projecting from selected 28 areas. No difference is detected in the summarized number of neurons projecting to LHA in WT and mSOD1 from the ipsilateral hemisphere (n=3). C: No difference is detected in the quantification of the absolute number of neurons projecting to LHA from 28 brain structures in WT and mSOD1. D: No difference is detected in the quantification of the normalized number of neurons projecting to LHA from 28 brain structures in WT and mSOD1. Bars show mean ± SD. Scalebars 1 mm.

***Supplementary Fig.7***: Unaltered cortico-hypothalamic projection pattern in mSOD1 mice at P25, contralateral hemisphere. A: Sum of neurons projecting from selected 28 areas. No difference is detected in the summarized number of neurons projecting to LHA in WT and mSOD1 from the contralateral hemisphere (n=3). B: Quantification of the absolute number of neurons projecting to LHA from 28 brain structures in WT and mSOD1. A significant increase in projections from ACA (p=0.0078) is detected. C: Quantification of the normalized number of neurons projecting to LHA from 28 brain structures in WT and mSOD1. No difference is detected in the quantification of the number of neurons projecting to LHA from 28 brain structures in WT and mSOD1. D: Representative *WholeBrain* volume reconstructions of neurons projecting to LHA in WT and mSOD1 mice show a similar pattern of projections in MOs (blue), ORBl/vl+AI (yellow) and PL+AI (green). Bars show mean ± SD. \**p* < 0.05, \*\**p* < 0.01, \*\*\**p* < 0.001, \*\*\*\**p*< 0.0001.

***Supplementary Fig.8***: Altered cortico-hypothalamic projection pattern in kiFUS mice at P270, absolute counts ipsilateral. A: Sum of neurons projecting from selected 28 areas. No difference is detected in the summarized number of neurons projecting to LHA in WT and kiFUS from the ipsilateral hemisphere (n=4). B: Quantification of the absolute number of neurons projecting to LHA from 28 brain structures in WT and mSOD1. A significant increase in projections from ACA (p=0.0385) and MOs (p<0.0001) is detected. C: Representative *WholeBrain* ipsilateral side view reconstructions of neurons projecting to LHA in WT and mSOD1 mice. Decrease of projections from MOs are visible (arrows). D: Representative *WholeBrain* cortical top view reconstructions of neurons projecting to LHA in WT and mSOD1 mice. Decrease of projections from MOs are visible (arrows). Bars show mean ± SD. \**p* < 0.05, \*\**p* < 0.01, \*\*\**p* < 0.001, \*\*\*\**p*< 0.0001.

***Supplementary Fig.9***: Altered cortico-hypothalamic projection pattern in kiFUS mice at P270, contralateral hemisphere. A: Sum of neurons projecting from selected 28 areas. No difference is detected in the summarized number of neurons projecting to LHA in WT and kiFUS from the contralateral hemisphere (n=3). B: Quantification of the absolute number of neurons projecting to LHA from 28 brain structures in WT and mSOD1. A significant increase in projections from ACA (p=0.0121) and ILA (p=0.0474) is detected. C: Quantification of the normalized number of neurons projecting to LHA from 28 brain structures in WT and kiFUS. A significant decrease in projections from ACA (p=0.0137) is detected. Bars show mean ± SD. \**p* < 0.05, \*\**p* < 0.01, \*\*\**p* < 0.001, \*\*\*\**p*< 0.0001.

## Notes

### Competing Interest Statement

The authors have declared no competing interest.

### Summary of Updates

We have updated and improved the DTI analysis section and the corresponding figure 5

