## Supplementary figures and images for "Orbitofrontal-hypothalamic projections are disrupted in hypermetabolic murine ALS model and human patients"

### Supplementary figure 1-9

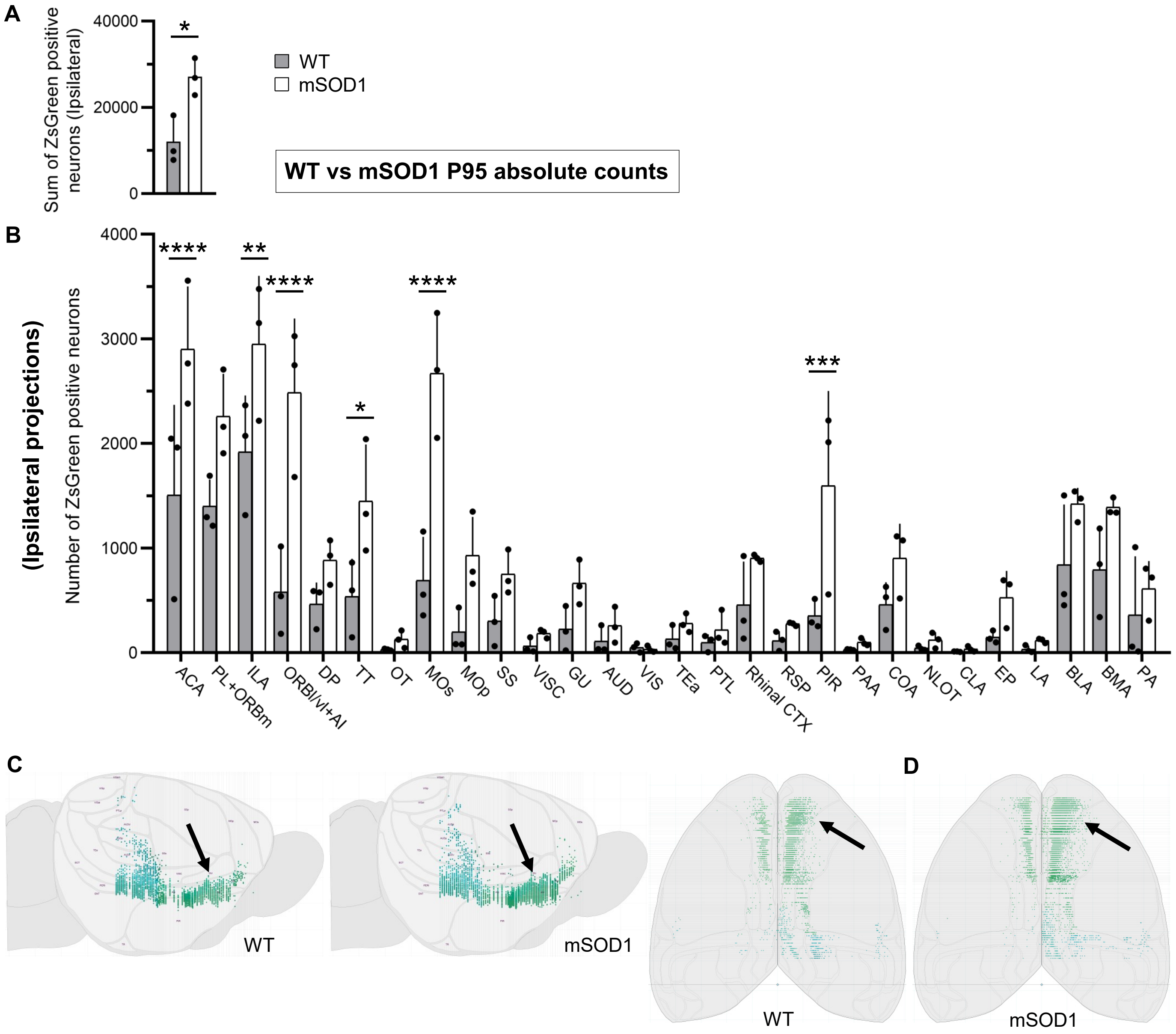

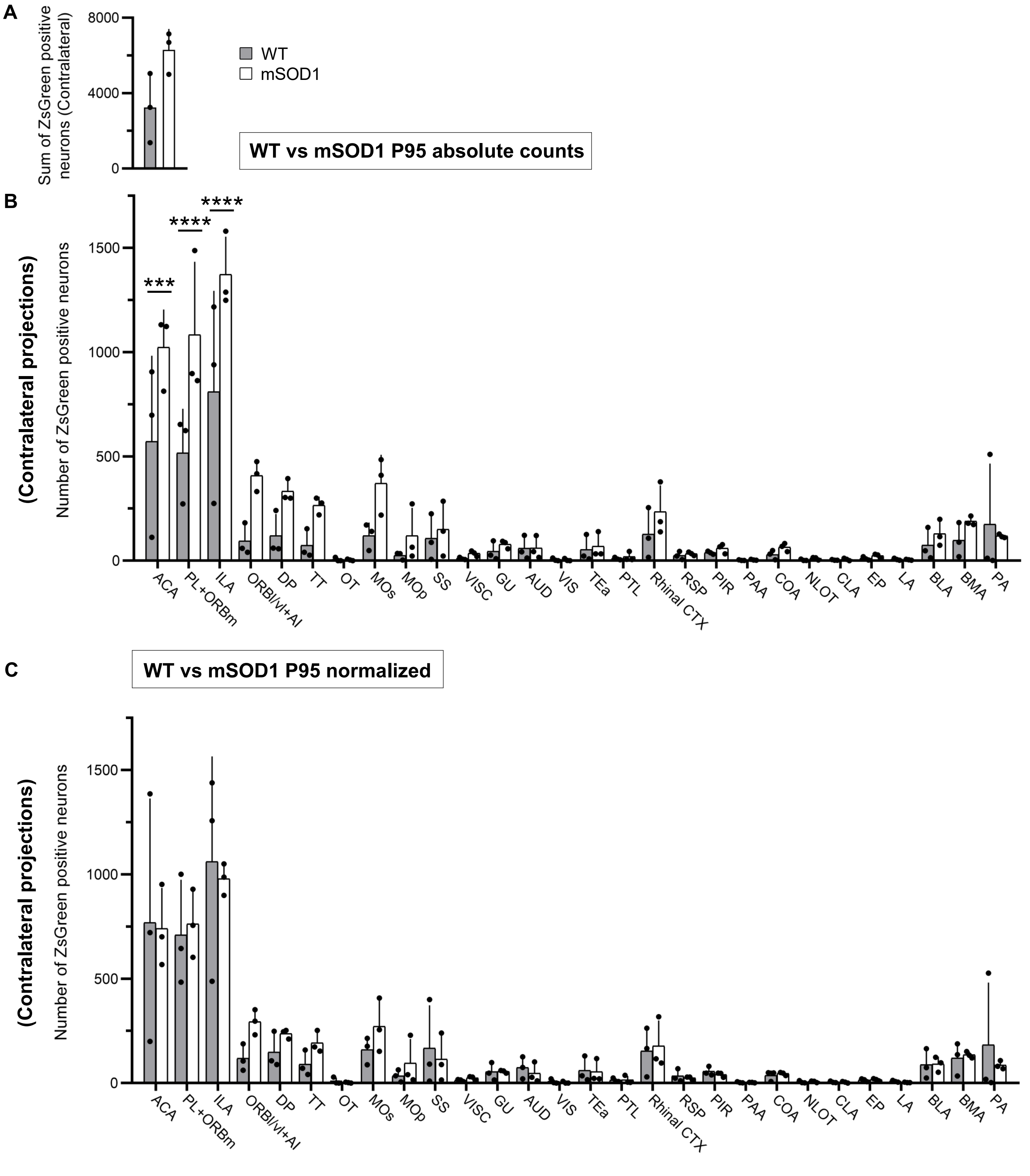

**WT vs mSOD1 P95**

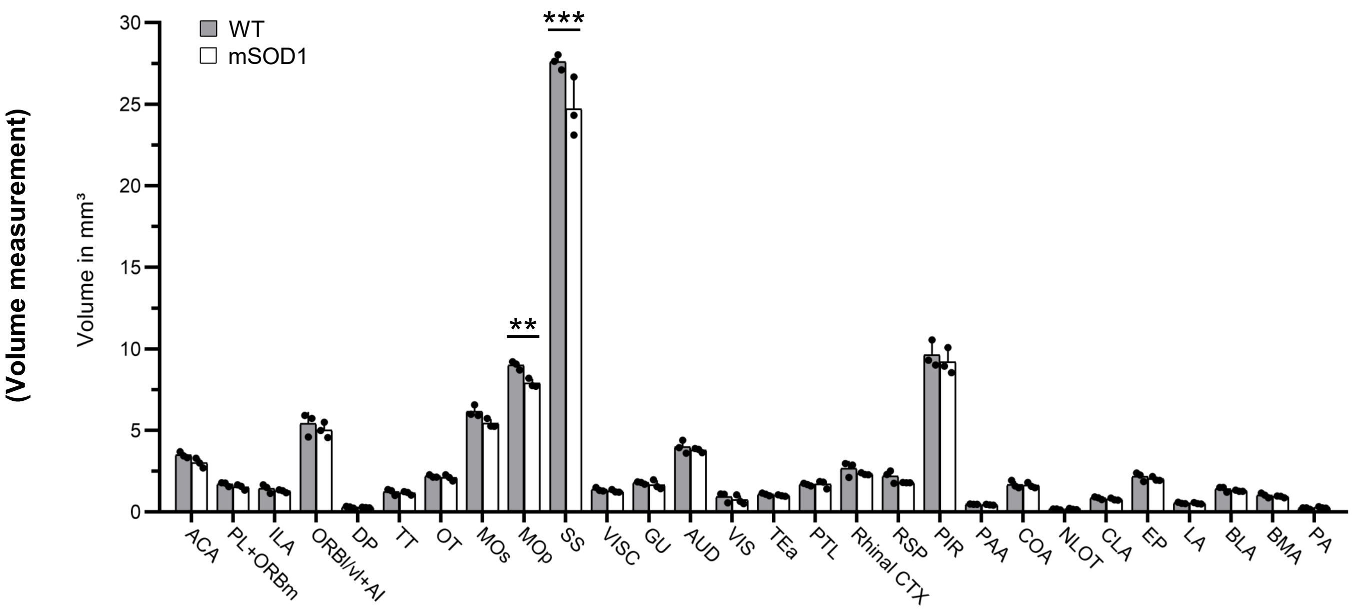

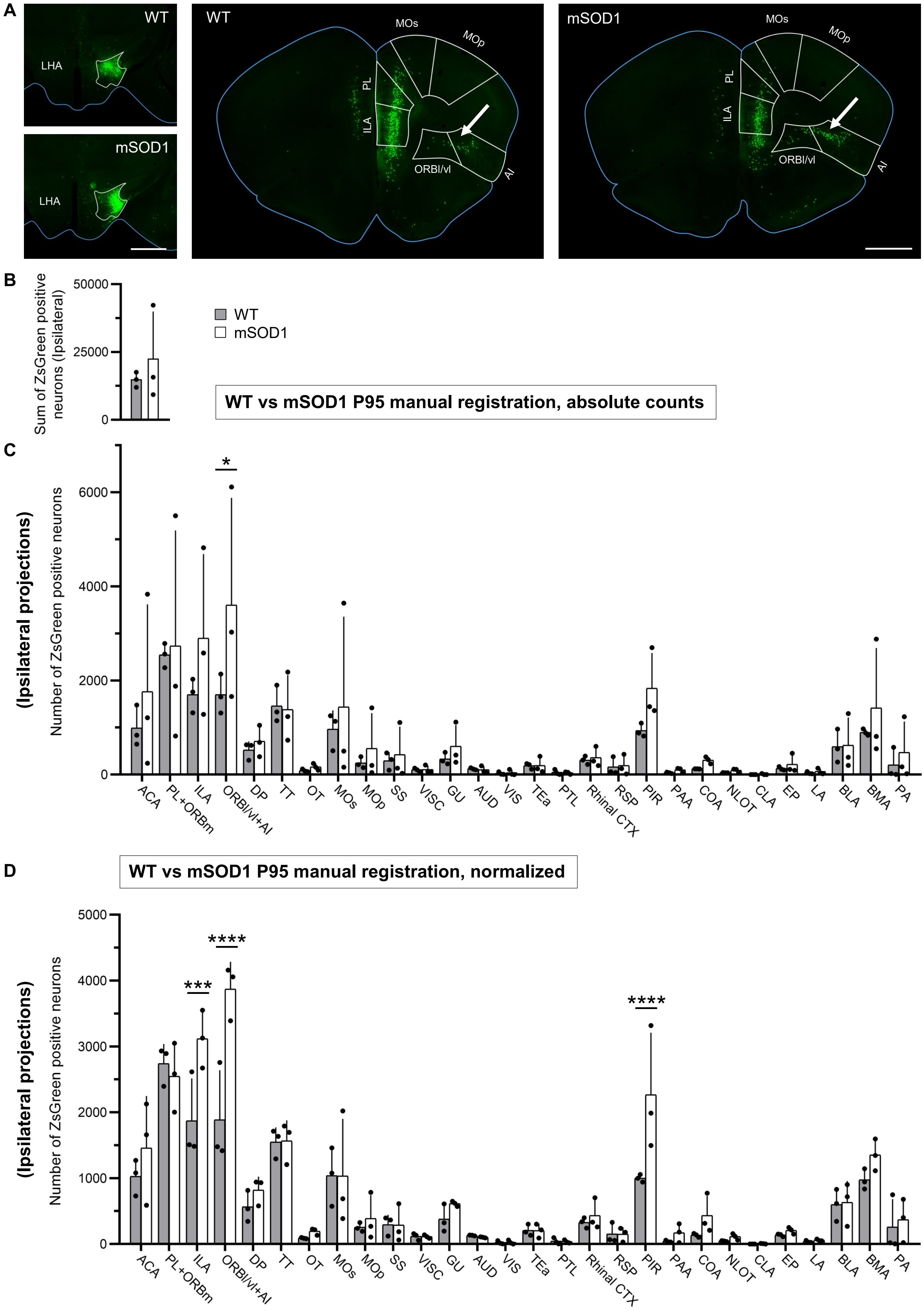

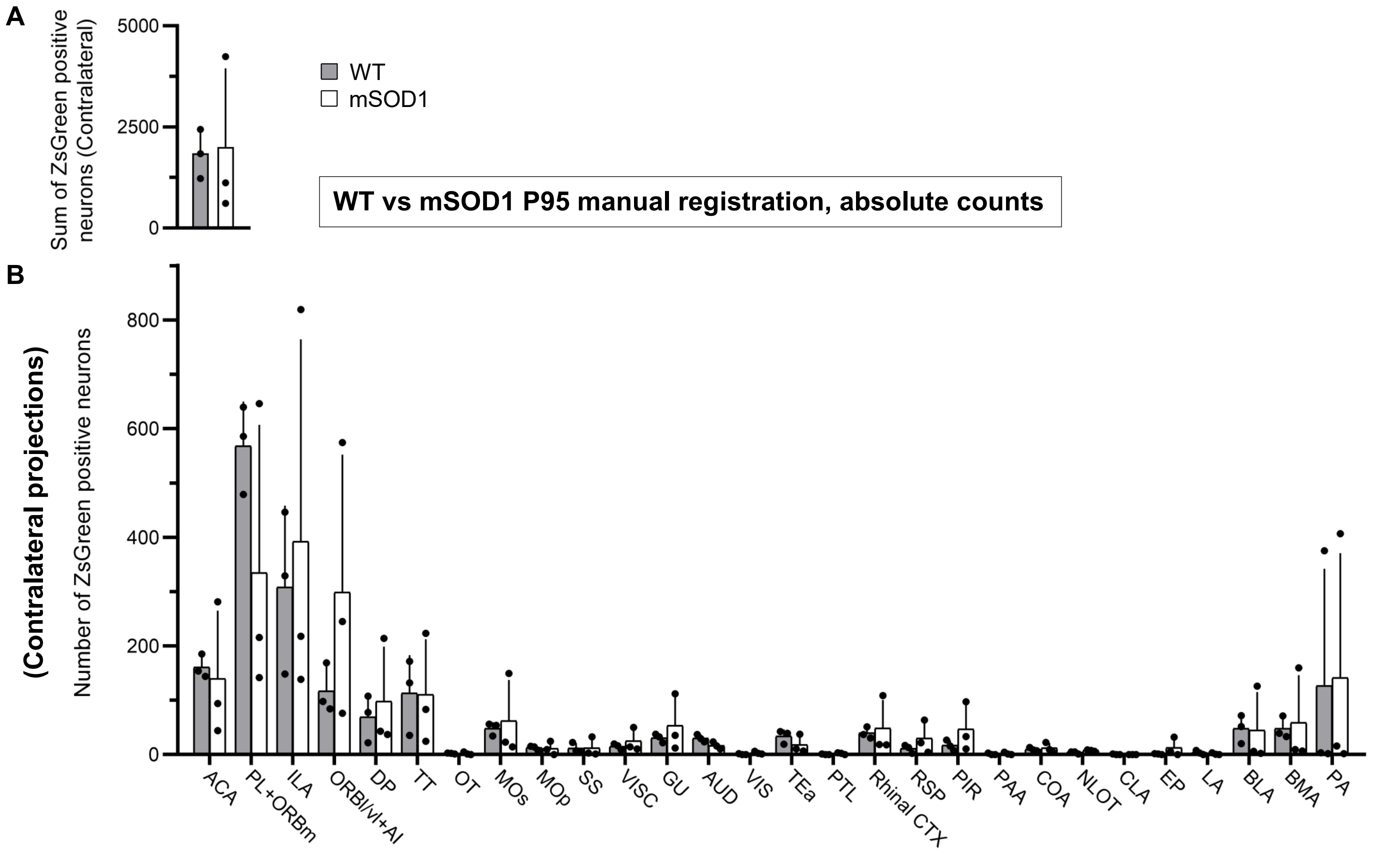

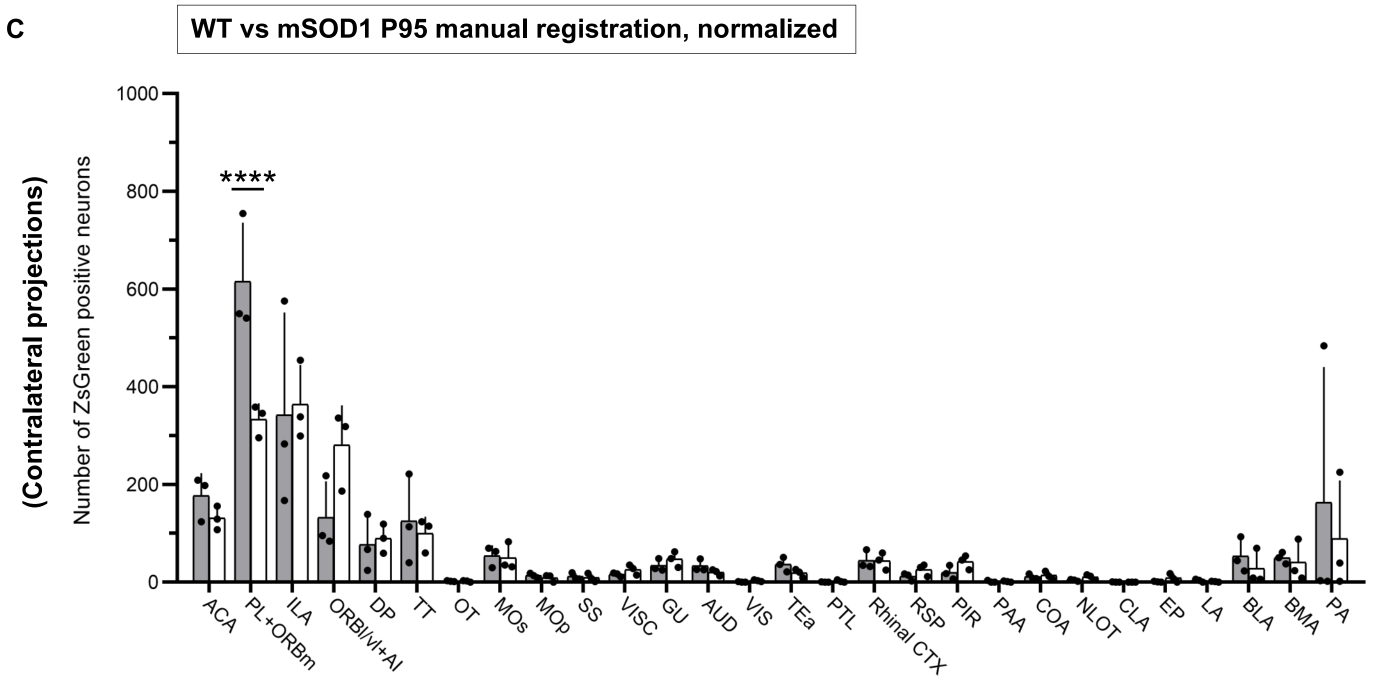

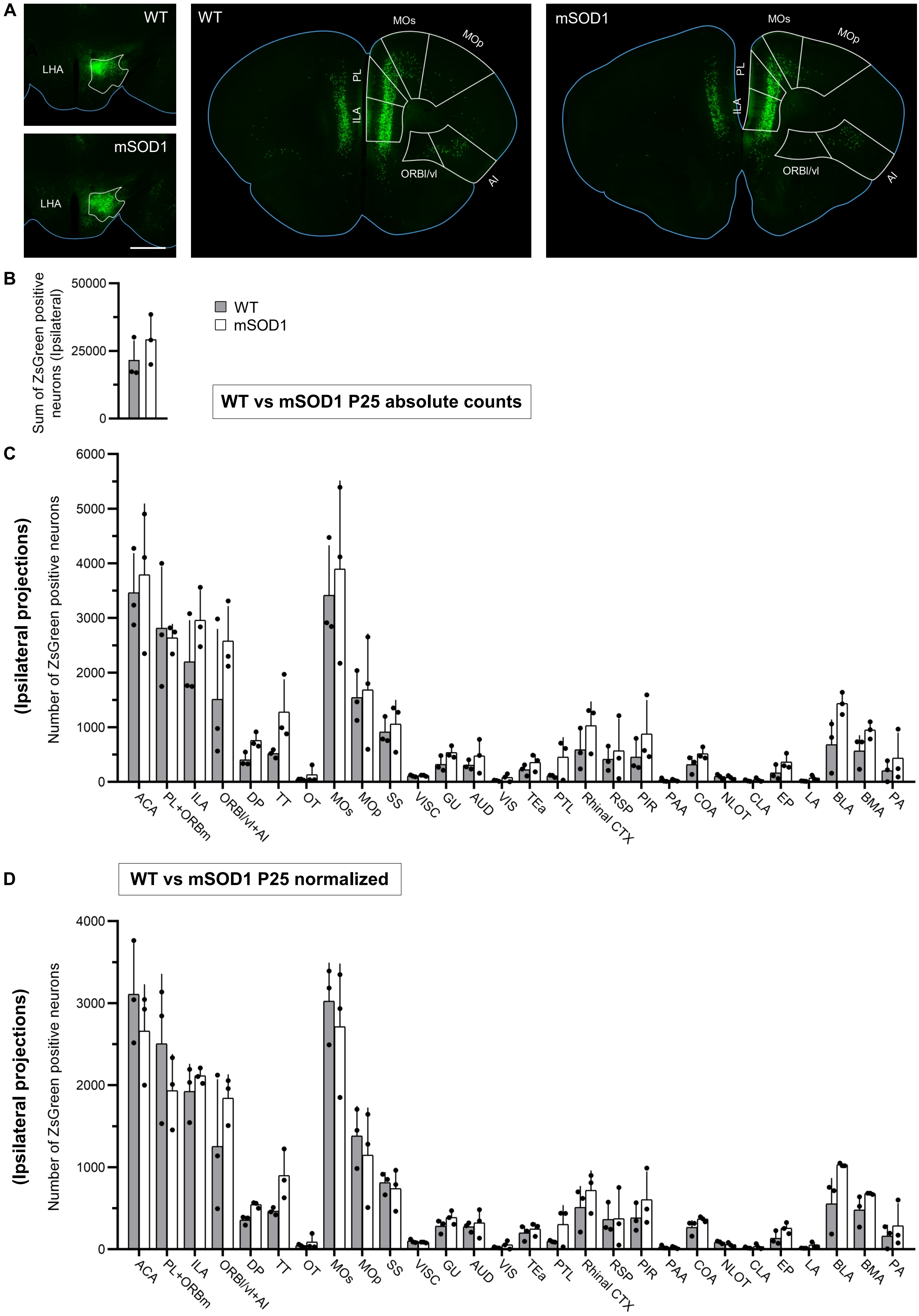

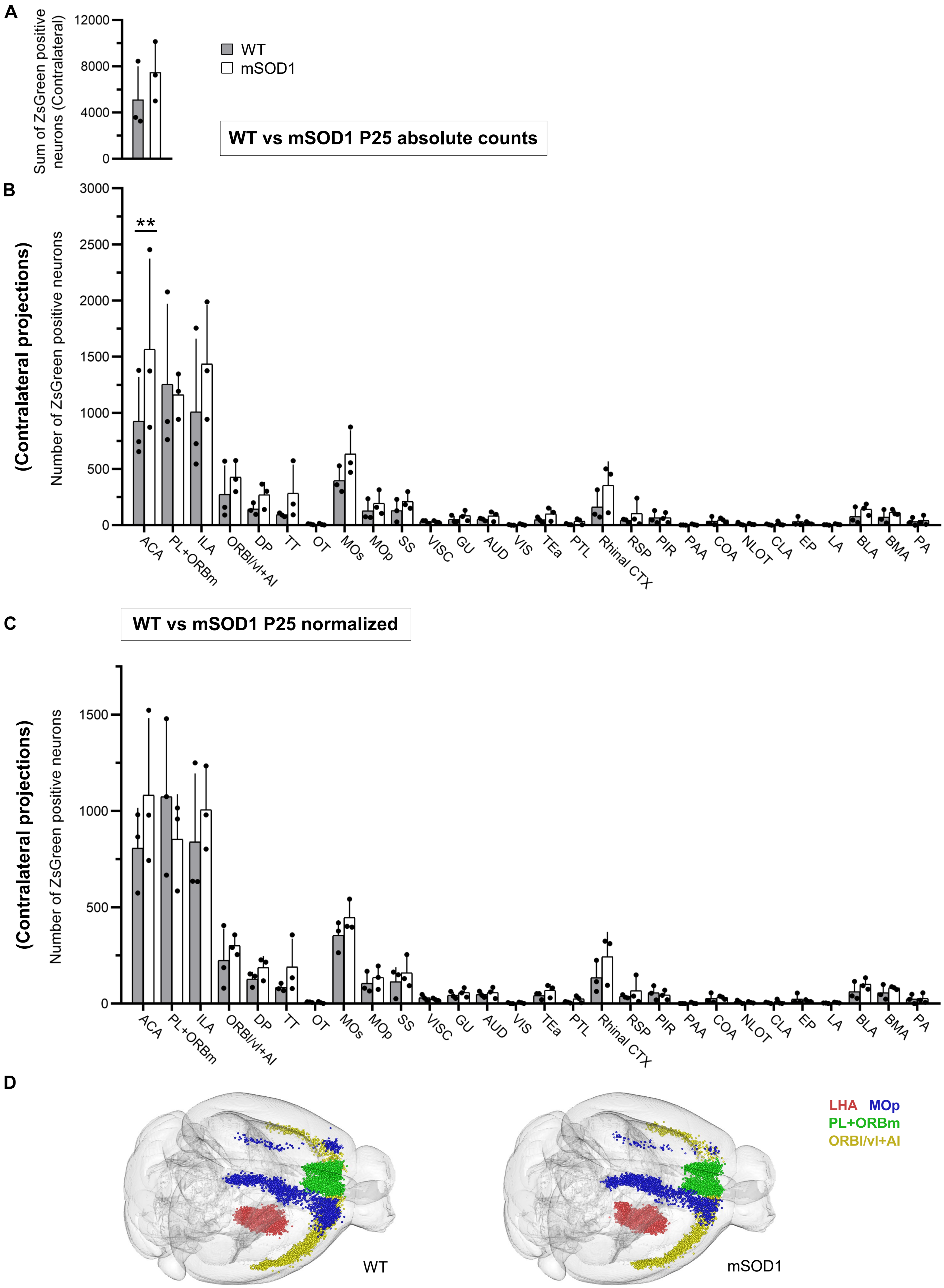

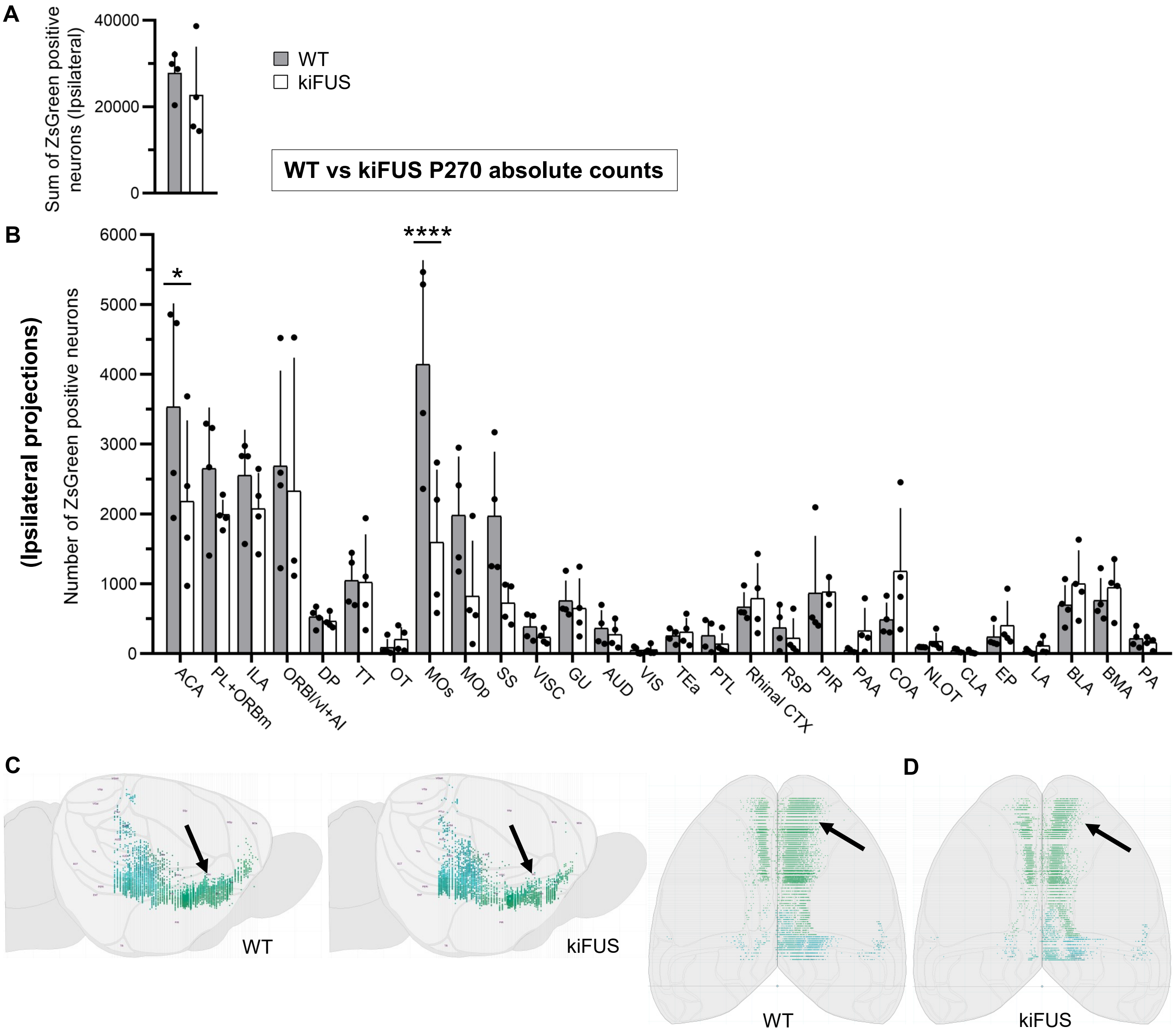

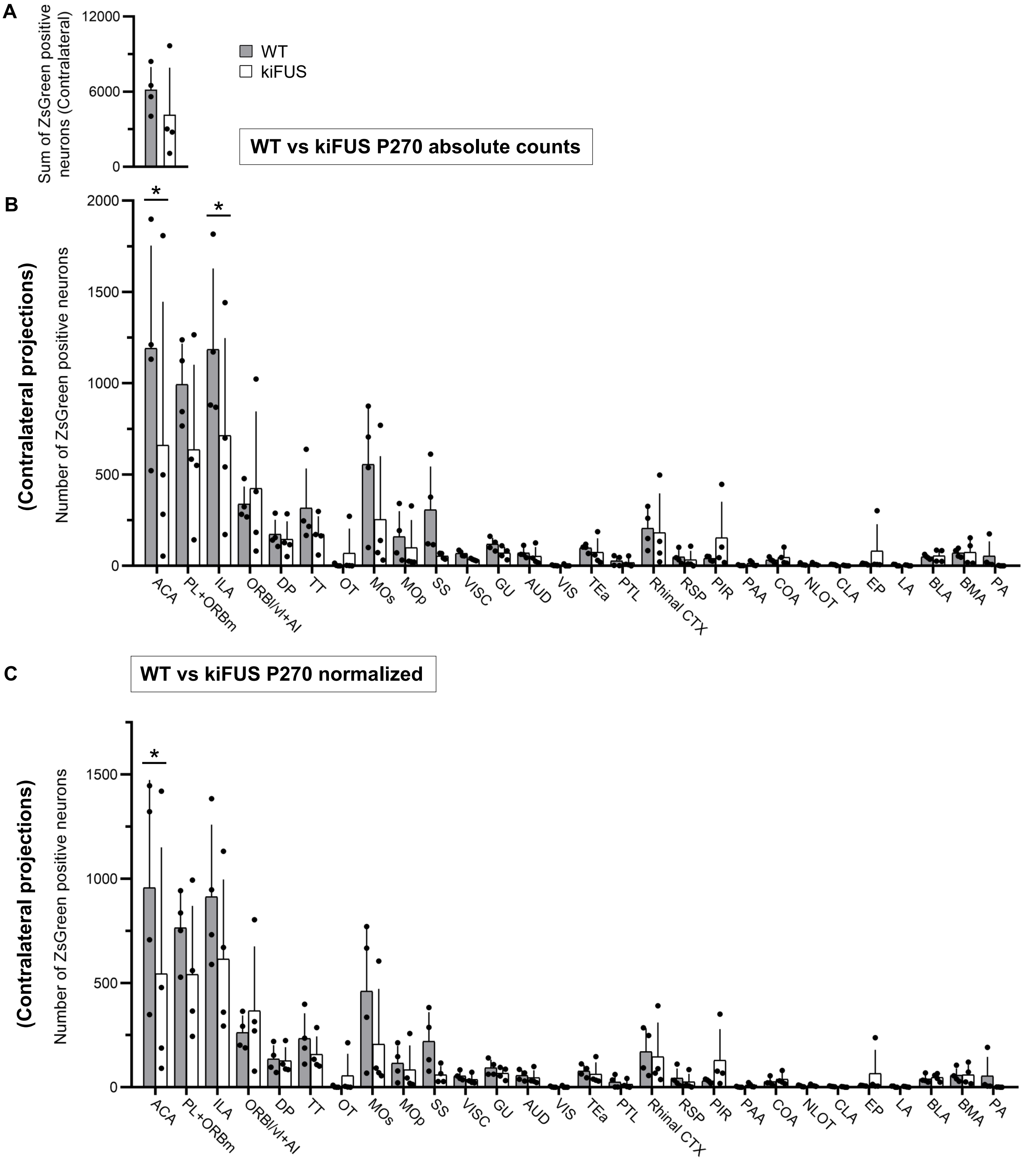
